# Matrix-M® adjuvant triggers inflammasome activation and enables antigen cross-presentation through induction of lysosomal membrane permeabilization

**DOI:** 10.1101/2025.04.30.651498

**Authors:** Behdad Zarnegar, Berit Carow, Jens Eriksson, Eva Spennare, Pontus Öhlund, Eray Akpinar, Emelie Bringeland, Ingrid Lekberg Osterman, Lena Lundqvist, Johanna Antti, Niklas Handin, Per-Henrik Helgesson, Johan Bankefors, Karin Lövgren Bengtsson, Mikael E. Sellin, Anna-Karin E. Palm, Linda Stertman, Carolina Lunderius Andersson

## Abstract

Matrix-M^®^ adjuvant, containing saponins, delivers a potent adjuvant effect and good safety profile. Given that Matrix-M is composed of Matrix-A and Matrix-C particles, comprising different saponin fractions, understanding their distinct roles can provide deeper insight into the mechanism of action of Matrix-M and guide future applications. Here, we demonstrate that antigen and Matrix-M, Matrix-A, or Matrix-C colocalize in lysosomes following uptake by bone marrow-derived dendritic cells. Matrix-M, Matrix-A, and Matrix-C induce lysosomal membrane permeabilization (LMP), but Matrix-C shows the highest LMP potential. LMP is required for interleukin (IL)-1β and IL-18 secretion *in vitro*. *In vivo*, a robust adjuvant effect of Matrix-M, Matrix-A, and Matrix-C is observed, both in the presence and absence of the NLRP3 inflammasome. LMP induced by Matrix-M, as well as Matrix-A and Matrix-C, also enables antigen cross-presentation. Thus, Matrix-induced LMP explains the capability of Matrix-M– adjuvanted protein vaccines to induce CD8^+^ T-cell responses.

## Introduction

Matrix-M® adjuvant is a component of COVID-19 vaccines that are approved or authorized for emergency use worldwide, and of malaria vaccines that are approved for use in Ghana, Nigeria, and Burkina Faso. These vaccines are well tolerated and have high efficacy for the prevention of COVID-19 and malaria, respectively, as found in phase 2/3 clinical trials^1–6^. Clinical studies have also shown that Matrix-M–adjuvanted vaccines induce a robust adaptive immune response, characterized by high levels of protective antibodies with broad epitope specificity, as well as antigen-specific T cells. The T-cell response includes T helper (Th)-1 biased CD4^+^ T cells, circulating follicular Th cells, and CD8^+^ T cells^1,7–10^.

Matrix-M adjuvant is a saponin-based adjuvant (SBA) derived from *Quillaja* saponins, which are extracted from the bark of the *Quillaja saponaria* Molina tree^11^. The bark extract is fractionated and fractionated saponins are formulated with phosphatidylcholine and cholesterol into 40-nm cage-like Matrix particles. Matrix-M contains two distinct adjuvant-active nanoparticles: Matrix-A particles with Fraction-A saponins and Matrix-C particles with Fraction-C saponins^11^. The mode of action of Matrix-M involves the rapid and transient presence of Matrix-M and activated innate immune cells at the injection site and its draining lymph nodes^12,13^. However, intracellular events following the uptake of Matrix-M and the molecular and cellular basis for its adjuvant effect have remained largely unknown.

The uptake of other SBAs, such as QS-21 and ISCOMATRIX™, into lysosomes has been shown to induce lysosomal membrane permeabilization (LMP), leading to the release of lysosomal contents into the cytosol^14–16^. This process results in downstream effects, including inflammasome activation and antigen cross-presentation. Lysosomes are single-membrane organelles that play a key role in cellular waste processing. They fuse with endosomes, phagosomes, and autophagosomes to degrade their contents, generating recycled components that can be used by the cell in various processes. This degradation is facilitated by hydrolytic enzymes, including proteases and lipases, that function optimally in the tightly regulated acidic environment of the lysosome^17^.

Inflammasomes, sensors of organelle damage or dysfunction, are activated following LMP^18–20^. Several SBAs have been reported to activate the NACHT, leucine-rich repeat, and pyrin domain domains containing protein 3 (NLRP3) inflammasome in innate immune cells^21,22^. Once activated, the inflammasome promotes the processing and secretion of the proinflammatory cytokines interleukin (IL)-1β and IL-18. This secretion is beneficial from an adjuvanticity perspective, as IL-1β supports the maturation of antigen-presenting cells (APCs) through enhancing lysosomal activity, upregulating the expression of major histocompatibility complex (MHC) and co-stimulatory molecules, and promoting the release of T-cell-activating cytokines^18^. IL-1 receptor signaling plays a key role in expanding naïve and memory T cells and prolonging T-cell help to B cells for antibody production. Additionally, the IL-1 family cytokine IL-18, which has been shown to contribute to ISCOMATRIX-mediated immunity, can be secreted in both an inflammasome-dependent and -independent manner^22^. IL-18, together with IL-12, supports the production of IFN-γ by NK cells, thereby promoting the differentiation of CD4^+^ T cells into Th1 cells. This mechanism is crucial for the induction of multifunctional Th1 cell responses, as seen with AS01, a liposome-formulated adjuvant containing QS-21 and the Toll-like receptor (TLR)-4 agonist 3-*O*-desacyl-monophorphoryl lipid A (MPL)^23^.

Protein-based vaccines are poor inducers of CD8^+^ T-cell responses, as exogenous antigens are typically presented by APCs via MHC class II (MHCII) molecules to prime CD4^+^ T cells. In order to activate antigen-specific CD8^+^ T cells, exogenous protein antigens need to enter the MHCI pathway. This process, known as antigen cross-presentation, requires the presence of a cross-presentation–inducing adjuvant as demonstrated for the SBAs ISCOMATRIX and Matrix-C^15,24–27^. Although Matrix-M–adjuvanted vaccines were reported to generate antigen-specific CD8^+^ T-cell responses in both mice and COVID-19 vaccine recipients, the mechanism of Matrix-M–induced antigen cross-presentation has not been examined^10,28,29^.

In the present study, we investigated the role of LMP in the adjuvant effect of Matrix-M and of its two constituent subcomponents Matrix-A and Matrix-C. Understanding the distinct functions of Matrix-A and Matrix-C will not only enhance our knowledge of the Matrix-M mechanism of action but may also allow for optimization and balancing of these components for targeted novel applications. Specifically, we examined the lysosomal localization of antigen and Matrix-M, Matrix-A, or Matrix-C after uptake by bone marrow-derived dendritic cells (BMDCs). We explored the distinct potential of Matrix-M, Matrix-A, and Matrix-C to induce LMP, which could lead to the release of lysosomal content into the cytosol. As downstream effects of LMP, we investigated the mechanisms underlying Matrix-induced NLRP3 inflammasome activation and its role in the adjuvant effect of Matrix-M. Additionally, we studied the ability of Matrix-M, Matrix-A, and Matrix-C to induce antigen cross-presentation and CD8^+^ T-cell activation in both *in vitro* and *in vivo* models.

## Materials and methods

### Adjuvants and antigens

Matrix-M, Matrix-A, and Matrix-C were obtained from Novavax AB. To enable cellular tracking, Matrix-A and Matrix-C were fluorescently labeled with BODIPY (4,4-difluoro-4-bora-3a,4a-diaza-s-indacen). The labeling process involved a HATU-mediated amide coupling reaction, coupling the BODIPY-FL-amine fluorophore (Lumiprobe GmbH) with the glucuronic acid unit within the branched trisaccharide domain of Fraction-A and Fraction-C saponins. Following synthesis, the labeled conjugates were purified using reverse-phase preparative high-performance liquid chromatography (RP-HPLC) on a C18 column and isolated as powders. The labeled particles were formulated using 90% unlabeled and 10% BODIPY-labeled Fraction-A or Fraction-C saponins for Matrix-A and Matrix-C, respectively. The final BODIPY-labeled Matrix-M consisted of 85% BODIPY-labeled Matrix-A and 15% BODIPY-labeled Matrix-C. For antigen studies, ovalbumin (OVA) protein (Endograde OVA, Lionex GmbH), the SIINFEKL peptide of OVA (Iba Lifesciences), and SARS-CoV-2 recombinant spike (rS) protein (BV2373, Novavax Inc.), which is formulated as a nanoparticle^29^, were used. For colocalization experiments, the SARS-CoV-2 rS was conjugated with Alexa Fluor™ 647 (Thermo Fisher Scientific) using NHS ester chemistry, followed by purification using Pierce™ dye removal columns (Thermo Fisher Scientific).

### Mice

Female BALB/c, C57BL/6, B6.129S6-*Nlrp3^tm1Bhk^*/J mice (referred to as *Nlrp3* knockout (KO) hereafter), and OT-I mice were obtained from Charles River Laboratories. All animal experiments were approved by the local animal experimentation ethical review committee (Uppsala djurförsöksetiska nämnd, approval number 5.8.18-12107/2021).

### Immunization

Mice (BALB/c mice for the Matrix dose titration; C57BL/6 and *Nlrp3* KO mice for the inflammasome study), 6‒10 weeks old, were immunized subcutaneously (s.c.) at the base of tail at day 0 and day 21 with 0.1 µg SARS-CoV-2 rS protein, either alone or adjuvanted with Matrix-M, Matrix-A, or Matrix-C in a total volume of 100 μL. Serum samples were collected at days 20 and 28, and additionally, at day 13 for C57BL/6 and *Nlrp3* KO mice.

### BMDC generation and cell culture

Primary BMDCs were generated by culturing total bone marrow cells (BMCs) isolated from tibia and femur of C57BL/6 and *Nlrp3* KO mice (1–3 mice/culture) in complete Roswell Park Memorial Institute (RPMI)-1640 medium (containing 10% heat inactivated fetal bovine serum, 2 mM L-glutamine, 100 U/ml penicillin and streptomycin, and 0.05 mM 2-mercaptoethanol) supplemented with 20 ng/ml recombinant murine granulocyte-macrophage colony-stimulating factor (GM-CSF) for 7 days. A total of 4×10^6^ BMCs were seeded per Petri dish, with half the medium replaced at days 3 and 6 with fresh medium containing GM-CSF (40 ng/mL). At day 7, non-adherent and loosely adherent cells were harvested by gentle PBS washing.

B3Z cells (a gift from Professor Sebastian Springer, Constructor University, Bremen, Germany) were maintained in complete RPMI-1640 medium supplemented with 0.5 mg/mL hygromycin B.

### Spinning-disk confocal microscopy

For spinning-disk confocal live-cell imaging, BMDCs (7.5 × 10^5^ cells/well) were seeded into sterile 8-well chambered cover glass (Cellvis) in phenol-red free complete RPMI-1640 medium supplemented with 20 ng/mL GM-CSF. On the following day, cells were stained with 50 nM LysoTracker™ Red DND-99 (Life Technologies) for 20 min and washed, and 10 pre-bleaching and background corrected images were immediately acquired at 3-min intervals. After this initial image recording, cells were exposed to BODIPY-labeled Matrix-M, Matrix-A, or Matrix-C, with or without the Alexa Fluor 647–labeled SARS-CoV2 rS antigen, and images were captured at 3-min intervals for an additional 120 min in DIC (transmitted light), GFP (Ex Lumencor 475/34x, Em Chroma ET510/20m), LysoTracker Red (Ex Lumencor 555/28x, Em Chroma ET600/50m) and Alexa Fluor 647 (Ex Lumencor 648/20x, Em Chroma 89402m) channels confocally, with a Spectra-X light engine (Lumencor) providing excitation light. A Prime™ 95B 25mm camera (Photometrics) coupled to an X-Light V2 spinning disk module (Crest Optics) on an ECLIPSE Ti2 microscope (Nikon) using a 100×/1.42 NA Plan Apochromat objective (Nikon) was used for imaging, pixel size 113 nm. During time-lapse imaging, the microscope chamber was maintained at 37 °C in a 5% CO_2_ moisturized atmosphere.

To visualize LMP, BMDCs (1.25 × 10^5^ cells/well) were seeded as described above and subsequently loaded with 50 μg/mL Alexa Fluor 488-conjugated 3-kDa dextran and 50 μg/mL tetramethylrhodamine-conjugated 40-kDa dextran (Molecular Probes) overnight. After washing, cells were chased in dextran-free medium for 2‒5 h to label lysosomes and incubated with or without 30 nM Bafilomycin A1 (BafA1, InvivoGen) for 1 hour. Cells were then imaged before and after treatment with Matrix-M, Matrix-A, or Matrix-C at 3-min intervals for up to 5 h to monitor the release of dextran from lysosomes to cytosol.

### Image analysis

The time-lapse movies were preprocessed by concatenating the pre- and post-treatment image series from individual positions, followed by subtraction of the background fluorescence from each fluorescence channel. For Matrix and antigen channels, backgrounds were generated by median projection of the first 10 pre-bleaching frames from each position. The LysoTracker Red background was generated by median projection across all positions and timepoints for each individual experiment. Pearson’s correlation coefficients (PCCs) for each time point and channel combination were calculated using the background-subtracted full frames by a Python (3.11.4) script using the scikit-image (0.22.0) and numpy (1.26.0) packages. The csv files output from the analysis script were further processed in R (3.5.1) and dplyr (1.0.5) and visualized with ggplot2 (3.3.3).

### Assessment of LMP by microscopy

One technical replicate (of three) from each experiment was analyzed. Prior to scoring all movies were pseudonymized and hence analysis was performed on blinded data to avoid scoring bias. Scoring was done manually using the point selection tool in ImageJ. Constant and global contrast settings were used, and cells were marked when diffuse dextran fluorescence was first detectable in the cytosol. All channels were independently analyzed in random order. The transmitted light channel (DIC) analysis comprised marking of when cells underwent lysis. All cells present in a given field of view were included in the analysis.

### Assessment of LMP by flow cytometry

BMDCs (1.5 × 10^5^ cells/well) were seeded in 96-well plates in complete RPMI-1640 medium supplemented with 20 ng/mL GM-CSF, and treated with serial dilutions of Matrix-M, Matrix-A, Matrix-C or medium for 1 h. Cells were then washed and resuspended using serum-free RPMI-1640 medium followed by staining with 50 nM LysoTracker™ Deep Red (Molecular Probes) and BD Horizon™ Fixable Viability Stain 520 (FVS520; BD Biosciences) for a total of 15 and 7 min, respectively. After washing, stained cells were detached using StemPro™ Accutase™ Cell Dissociation Reagent (Gibco) and analyzed on a BD FACSCelesta™ flow cytometer (BD Biosciences). Data were processed using FlowJo software version 10.7.1 (BD Life Sciences). As positive control for reduced LysoTracker staining, cells treated with BafA1 for 1 h or with *L*-Leucyl-*L*-Leucine methyl ester (LLOMe, Merck) for 15 min, were analyzed in parallel. Normalized geometric means of fluorescence intensity (gMFI) for LysoTracker were calculated relative to untreated controls.

### Assessment of inflammasome activation, cell viability and cytokine production

BMDCs were seeded in 96-well plates for viability assays (0.5 × 10^5^ cells/well) and for cytokine analysis by ELISA (1 × 10^5^ cells/well). Cells were either left unprimed or primed with 0.1 µg/mL LPS (Merck) for 1 h, followed by treatment with MCC950, BafA1, Ac-YVAD-cmk (all from InvivoGen), CA074-Me (Selleck Chemicals), and disulfiram or necrosulfamide (Tocris Bioscience) for an additional 1 hour. Cells were then exposed to serial dilutions of Matrix-M, Matrix-A, or Matrix-C. After 5 or 24 h of incubation with Matrix, cell viability was assessed, and supernatants were harvested and immediately frozen for subsequent cytokine analysis by ELISA. All treatments were performed in duplicate wells. Cell viability was determined using WST-1 cell proliferation reagent (Roche), and supernatants were analyzed for IL-1β and IL-18 using mouse IL-1β and IL-18 ELISA kits according to the manufacturer’s instructions (DuoSet ELISA, R&D Systems).

### Antigen cross-presentation and BMDC phenotyping *in vitro*

BMDCs (1.5 × 10^5^ cells/well) were seeded in non-treated 96-well round-bottom plates and incubated for 24 h with 0.8 mg/mL OVA (EndoGrade®Ovalbumin, Lionex GmbH) and Matrix-M, Matrix-A, or Matrix-C. For flow cytometry analysis, cells were stained with FVS780 (BD Biosciences). After washing, cells were detached with PBS-EDTA (Alfa Aesar, Thermo Fisher Scientific), resuspended in FBS-containing stain buffer (BD Biosciences), incubated with an anti-mouse CD16/CD32 antibody (2.4G2), followed by staining with F4/80:BV605 (T45-2342), CD11c:BV650 (HL3), I-E/I-E(MHCII):BV786 (M5/114.15.2), CD86:PE (GL1), CD11b:PerCP-Cy5.5 (M1/70), Ly6C:R718 (AL21, all BD Biosciences), and SIINFEKL-H-2kB (25-D1.16; eBioscience) for 20 min at 4 °C in the dark. Stained cells were washed and immediately analyzed on a BD FACSCelesta flow cytometer with data analysis performed using FlowJo software version 10.9.0.

### B3Z assay

BMDCs (8 × 10^4^ cells/well) were seeded in 96-well flat-bottom cell culture plates in the presence or absence of BafA1 or epoxomicin for 1 h and incubated with 80 µg/mL OVA and Matrix-M, Matrix-A, or Matrix-C for 5 h. As a control for cell viability or MHCI expression levels, control BMDCs (without OVA) were pulsed with 5 ng/mL SIINFEKL peptide 30 min before the incubation ended. Cells were washed twice with PBS and co-cultured with B3Z cells (8 × 10^4^ cells/well) for 18 h, with BafA1 or epoxomicin in treated wells. Cells were then washed twice and lysed in a PBS buffer containing 0.15 mM CPRG (Roche), 9 mM MgCl_2_, 0.1 mM 2-mercaptoethanol, and 0.125% NP-40. CPRG conversion by β-galactosidase was measured after 3–5 h at 570 nm using a SpectraMax® M3 spectrophotometer (Molecular Devices).

### Antigen cross-presentation *ex vivo*

Female C57BL/6 mice (8–10 weeks) were immunized with 50 µL intramuscular (i.m.) injection of OVA, either alone or adjuvanted with Matrix-M, into the right hind leg (*musculus quadriceps femoris*). The draining iliac lymph nodes (LN) were collected 24 h post-immunization and the obtained cell suspensions were pooled (7 to 13 LN per pool) and sorted by MACS using the CD11c MicroBeads Ultrapure (Miltenyi Biotec) to isolate CD11c^+^ dendritic cells (DCs). These CD11c^+^ DCs were co-cultured at a 1:4 ratio in a 96-well U-bottom cell culture plate with naïve CD8^+^ T cells isolated from spleens of OT-I mice using the Naïve CD8a^+^ T cell Isolation Kit (Miltenyi Biotec). OT-I cells had been pre-labeled with CellTrace™ Violet (CTV, Life technologies) to follow their proliferation. After three days of co-culture, cells were stained with FVS780, followed by incubation with anti-mouse CD16/CD32 antibody (2.4G2; Fc block) and antibodies targeting CD4:PerCyP5.5 (RM4-5), CD8a:BV650 (53-6.7), CD44:BV786 (IM7), and CD3e:FITC (145-2C11, all antibodies from BD Biosciences) for 30 min at 4 °C in the dark. Stained cells were washed and immediately analyzed on a BD FACSCelesta flow cytometer with data analysis performed using FlowJo software version 10.9.0. The MFI of CD44:BV786 for gated CD8^+^ T cells was used to assess their activation, and CTV dilution was analyzed to determine the percentage of divided CD8^+^ T cells using FlowJo’s proliferation modeling tool, annotating the “generation 0” peak for undivided cells and number of observed peaks.

### ELISA for anti-rS IgG1 and IgG2a/IgG2c serum antibodies

Quantification of anti-rS IgG1 and IgG2a or IgG2c serum antibodies (BALB/c or C57BL/6 background, respectively) was performed using ELISA as previously described^13^. Serum samples were serially diluted 5-fold in eight steps, starting at 1:15 or 1:50 (days 13 and 20), 1:500, 1:5000, or 1:10,000 (day 28) for IgG1, and 1:15 (day 20), 1:15, 1:100, or 1:500 (day 28) for IgG2a, and 1:15 or 1:50 (days 13 and 20), 1:15, 1:50, 1:100, or 1:500 (day 28) for IgG2c. Absorbance was measured at 450 nm (SpectraMax M3, Molecular Devices) and anti-rS titers were calculated using a four-parameter logistic equation (SoftMax® Pro software v.7.1.1.). Pooled sera from untreated or from SARS-CoV-2 rS (adjuvanted with Matrix-M) immunized mice were used as negative and positive controls, respectively.

### ELISA for hACE2 receptor-blocking antibodies

Quantification of hACE2 receptor-blocking antibody titers was performed in an ELISA assay as previously described^13^. Mouse sera were serially diluted 2-fold (starting with a 1:20 dilution). Absorbance was measured at 450 nm (SpectraMax M3, Molecular Devices). Serum antibody titers at 50% inhibition (IC50) of hACE2 to SARS-CoV-2 rS protein were then determined using the SoftMax Pro software v.7.1.1.

### Cellular responses using FluroSpot assay

Antigen-specific recall cellular responses were evaluated at day 28 via the FluoroSpot assay as previously described^13^. In short, single-cell suspensions were seeded on FluoroSpot filter plates (Mabtech) precoated with anti–IL-2, anti–IFN-γ, or anti–IL-4 capture antibodies. Cells were stimulated with 1 μg/well SARS-CoV-2 rS protein and after incubation cytokine-positive spots were developed according to the manufacturer’s instructions and counted using an iSpot FluoroSpot Reader Spectrum (Autoimmune Diagnostika GmbH). The number of IFN-γ– and/or IL-2–, or IL-4–secreting cells was obtained by subtracting the mean background number of the respective medium control for each mouse, followed by averaging triplicate wells. The data are shown as geometric mean (GM) for each group. The results are expressed as the number of cytokine-secreting cells per 10^6^ splenocytes.

### T-cell responses using Intracellular Cytokine Staining (ICS)

For ICS, splenocytes (1 × 10^6^ cells/well) were cultured in U-bottom 96-well cell culture plates and stimulated with 5 μg/mL SARS-CoV-2 rS overnight. Next, plates were incubated with GolgiStop™ and GolgiPlug™ (BD Biosciences) for 5 hours. Cells were stained with FVS780, washed and incubated with an anti-mouse CD16/CD32 antibody (2.4G2) antibodies targeting CD3:R718 (clone 145-2C11), CD4:PerCpCy5.5 (RM4-5), CD8a:BV650 (53-6.7), and CD44:BV786 (IM7) (BD Biosciences). Cells were incubated with BD Cytofix/Cytoperm™ Solution (BD Biosciences) for 20 min at 4 °C in the dark and kept in PBS overnight after washing. The next day, cells were incubated with Perm/Wash™ Buffer (BD Biosciences) for 15 min at room temperature and stained with IFN-γ:BV421 (clone XMG1.2), TNF:PE (MP6-XT22), IL-2:PE-Cy7 (JES6-5H4), and IL-4:APC (11B11)(BD Biosciences). After washing, stained cells were analyzed on a BD FACSCelesta flow cytometer and data processed using FlowJo software version 10.9.0. Live CD3^+^ T cells were gated for CD4^+^ and CD8^+^ T cells. Within CD4^+^ and CD8^+^ T cells, the frequency of single cytokine-positive/CD44^+^ cells was determined. Frequencies of unstimulated medium controls were subtracted from rS-stimulated matched samples for background reduction.

### Statistical analysis

Specific information on sample sizes, number of experiments, technical replicates and statistical analyses is provided in the figure legends. *p*-values of less than 0.05 were considered statistically significant (\**p* ≤ 0.05; \*\**p* ≤ 0.01; \*\*\**p* ≤ 0.001; \*\*\*\**p* ≤ 0.0001). The graphs were prepared, and statistics calculated using GraphPad Prism® version 9.4.0 or later (GraphPad Software).

## Results

### Matrix-M accumulates inside lysosomes

In order to determine the intracellular localization of Matrix-M after cellular uptake, we employed both BODIPY-labeled Matrix-M and LysoTracker. BODIPY labeling of Matrix-A and Matrix-C particles enabled us to observe the trafficking of Matrix-M within the cells, while LysoTracker, which selectively accumulates in acidic organelles such as lysosomes upon protonation, allowed assessment of whether Matrix-M localizes to lysosomes through the endocytic pathway^30^. For this, BMDCs were stained with LysoTracker, followed by incubation with BODIPY-conjugated Matrix-M. Using time-lapse spinning-disk confocal imaging, we observed that shortly after exposure, BODIPY-labeled Matrix-M became visible inside the cells (**Fig. 1A)** with a time- and dose-dependent increase in BODIPY fluorescent signal (**Fig. 1B**). Image overlays displayed a substantial overlap between LysoTracker and BODIPY signals (**Fig. 1A**). Furthermore, by calculating Pearson’s correlation coefficient (PCC) for each frame, a marked increase in the degree of colocalization between LysoTracker and BODIPY-labeled Matrix-M was observed (**Fig. 1C, Supplementary Fig. 1**). Similarly, cellular uptake and lysosomal localization were observed using BODIPY-labeled Matrix-A or BODIPY-labeled Matrix-C (**Fig. 1A–C**). Matrix-C displayed a higher fluorescence intensity compared to Matrix-A at the highest tested concentration (5 μg/mL) (**Fig. 1B**). By contrast, Matrix-A reached a higher level of colocalization with LysoTracker than Matrix-C (**Fig. 1C**). Altogether, these results indicate an accumulation of Matrix-M, Matrix-A, and Matrix-C within lysosomes upon cellular uptake.

**Figure 1:**
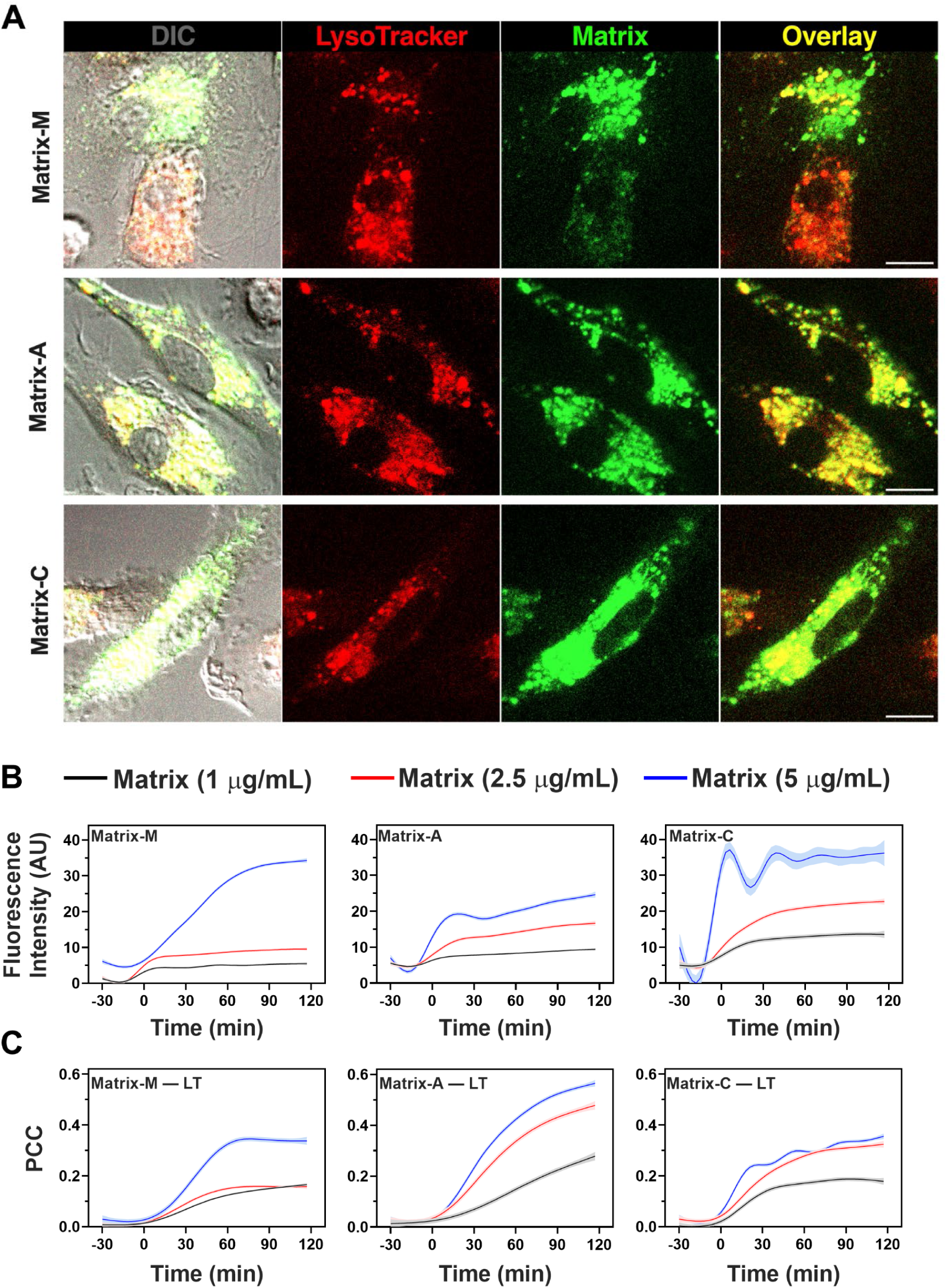
Matrix-M accumulates inside lysosomes. BMDCs were loaded with LysoTracker Red DND-99, washed and imaged for 30 min. Following the initial pre-bleaching image acquisition, cells were treated with BODIPY-labeled Matrix-M, Matrix-A, or Matrix-C at concentrations of 1, 2.5, or 5 μg/mL, and live-cell spinning-disk confocal imaging continued for an additional 120 minutes. **(A)** Representative images showing individual channels (LysoTracker, BODIPY), overlays, and the DIC (Differential Interference Contrast) channel captured after each treatment (scalebar = 10 μm). **(B)** The fluorescence intensity kinetics of BODIPY fluorescence for the indicated Matrix concentrations. **(C)** Pearson’s correlation coefficients (PCC) for BODIPY and LysoTracker fluorescence over time. Data shown in **B** and **C** represent smoothed averages from at least three technical and biological replicates ± SEM.

### Matrix-M induces lysosomal membrane permeabilization (LMP) leading to the release of lysosomal contents into the cytosol

To evaluate the effect of Matrix-M on lysosomal membrane integrity, which is required to maintain a low intra-lysosomal pH, we assessed lysosomal acidification by monitoring LysoTracker cellular staining using flow cytometry. In viable BMDCs (gating strategy shown in **Supplementary Fig. 2**), exposure to Matrix-M adjuvant resulted in a reduction of LysoTracker staining (**Fig. 2A, B**). Similar reductions in LysoTracker staining were observed following exposure to Matrix-A and Matrix-C, indicating that all three Matrix adjuvants disrupt the pH gradient across the lysosomal membrane, or disrupt the lysosomal membranes themselves. As expected, the positive controls, BafA1 and LLOMe, also reduced LysoTracker staining (**Fig. 2A, B**). BafA1 and LLOMe disrupt lysosomal acidification via distinct mechanisms. BafA1 inhibits the vacuolar ATPase (V-ATPase) proton pump responsible for maintaining low lysosomal pH^31^, whereas LLOMe induces LMP by forming membrane pores^32,33^. Visible Matrix-M– induced reduction of LysoTracker signal could also be noted among single cells imaged by spinning-disk confocal live-cell microscopy (**Fig. 2C**).

**Figure 2:**
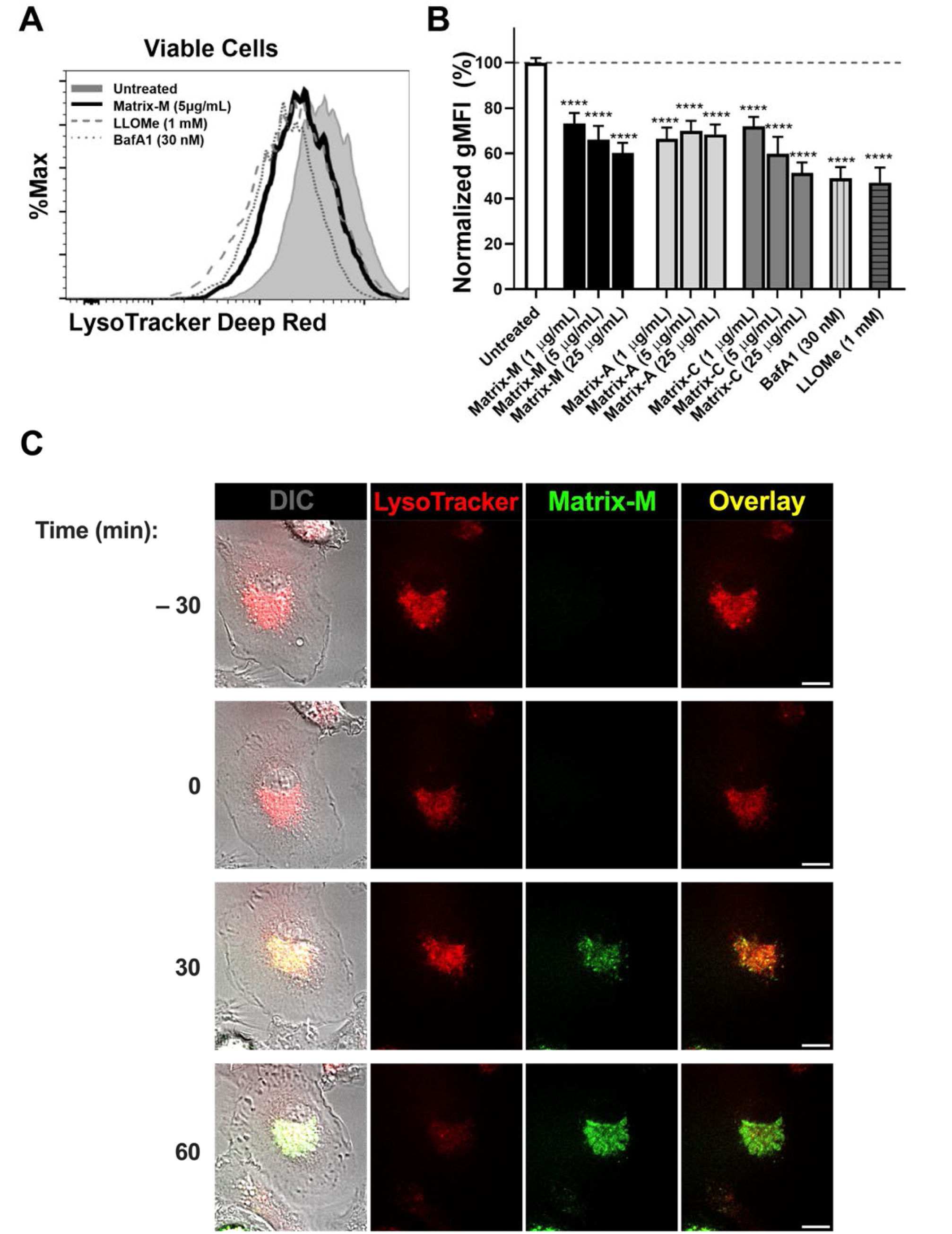
Matrix-M induces lysosomal membrane permeabilization (LMP) BMDCs were treated with either Matrix-M, Matrix-A, or Matrix-C (for 1 h), bafilomycin (BafA1, for 1 h), or *L*-Leucyl-*L*-Leucine methyl ester (LLOMe, for 15 min) at the indicated concentrations followed by staining with LysoTracker Deep Red and FVS520 for 15 and 7 min, respectively. Stained cells were analyzed by flow cytometry. **(A)** Flow cytometry histograms showing LysoTracker fluorescence intensity in viable BMDCs from representative samples (gating shown in **Supplementary Fig. 2**). **(B)** Normalized geometric mean fluorescence intensity (gMFI) of LysoTracker Deep Red in BMDCs. Data were normalized to untreated control samples and are shown as the mean ± SD; pooled from three independent experiments. Statistical differences between untreated control and treated samples were determined by one-way ANOVA followed by Dunnett’s multiple comparisons test (****p < 0.0001). **(C)** Images of LysoTracker-stained BMDCs before and after treatment with BODIPY-labeled Matrix-M, as described in Figure 1. Scalebar = 10 μm.

We next sought to investigate whether the Matrix-induced loss of lysosomal acidification could be attributed to pore-forming activity of Matrix-M, leading to LMP. To this end, we monitored induction of LMP by tracking the translocation of fluorescent dextran particles (3- and 40-kDa) from lysosomes to the cytosol using time-lapse spinning-disk confocal imaging^34^. Before the addition of Matrix-M, cells had intact lysosomes as indicated by the punctate morphology of the fluorescent dextran particles (**Fig. 3A**). However, a diffuse cytosolic staining pattern for both 3- and 40-kDa dextran particles appeared in the cells upon Matrix-M treatment, demonstrating the induction of pores permeable for particles up to 40-kDa size (**Fig. 3A**). Similarly, the treatment of dextran-loaded BMDCs with Matrix-A or Matrix-C resulted in the release of lysosomal content into the cytosol (**Fig. 3A**). Of note, Matrix-M and Matrix-C were found to be more potent than Matrix-A in promoting LMP and dextran release into the cytosol (**Fig. 3B**, note different concentrations of Matrix adjuvants). Given that lysosomal hydrolytic enzymes depend on acidic pH for optimal activity, we next assessed whether lysosomal acidification is required for the LMP-inducing effect of Matrix adjuvants. For this, BMDCs preloaded with dextran particles were incubated with the lysosomal acidification inhibitor BafA1 for 1 h before exposure to Matrix-M, Matrix-A, or Matrix-C. Notably, inhibition of lysosomal acidification completely blocked the release of both 3- and 40-kDa dextran particles from the lysosome to the cytosol in response to Matrix-M and Matrix-A (**Fig. 3B)**. In response to Matrix-C, blockade of lysosomal acidification resulted in complete inhibition of 40-kDa dextran release, but only partial inhibition of 3-kDa dextran release into the cytosol (**Fig 3B**). These results demonstrate that all three Matrix adjuvants induce LMP, albeit with distinct potencies, as Matrix-C and Matrix-M exhibited stronger LMP-inducing effects compared to Matrix-A.

**Figure 3:**
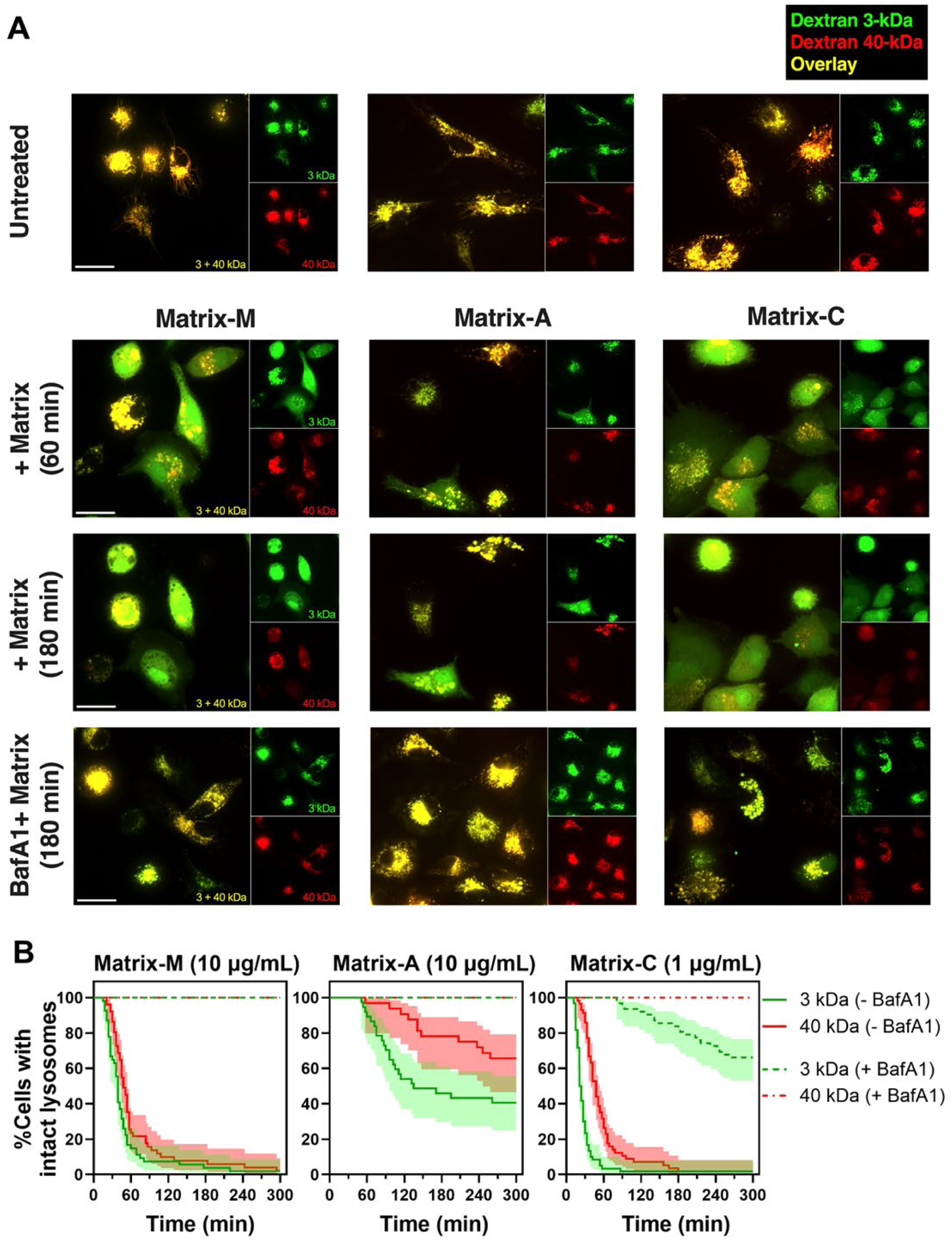
Matrix-induced LMP results in the release of lysosomal contents into the cytosol. BMDCs were preloaded overnight with dextran particles of different sizes and fluorochromes (Alexa Fluor 488-conjugated 3-kDa dextran and tetramethylrhodamine-conjugated 40-kDa dextran). The following day, cells were pretreated with bafilomycin A1 (BafA1, 30 nM) or left untreated for 1 hour. Live-cell spinning-disk confocal imaging was performed at 3 min intervals for up to 5 h, before and after adding Matrix-M, Matrix-A (both at 10 μg/ml), or Matrix-C (1 μg/ml), to monitor lysosomal release of dextran particles into the cytosol. **(A)** Representative images showing individual channels (3- and 40-kDa dextrans) and overlays at the indicated time points and treatment conditions (scalebar = 20 μm). **(B)** The percentage of cells with intact lysosomes over time for each treatment condition. Data are pooled from three independent experiments and presented as percent cells with intact lysosomes ± 95% confidence interval.

### Matrix-M induces inflammasome activation and IL-1β secretion

Release of lysosomal contents into the cytosol following LMP can initiate inflammasome activation, a process essential for converting pro–IL-1β into its active form and the subsequent release of biologically active IL-1β. To assess the ability of Matrix-M to induce inflammasome activation, non-primed and LPS-primed BMDCs were treated with increasing doses of Matrix-M for 5 h, and released IL-1β levels were measured in supernatants by ELISA. Matrix-M induced the release of IL-1β from LPS-primed BMDCs, with higher doses of Matrix-M resulting in greater IL-1β secretion (**Fig. 4A**). Similarly, a dose-dependent secretion of IL-1β was observed following exposure of the cells to Matrix-A or Matrix-C. Notably, Matrix-M induced higher levels of IL-1β than either Matrix-A or Matrix-C. By contrast, non-LPS primed BMDCs did not release IL-1β upon treatment with any of the Matrix adjuvants. To investigate whether lysosomal acidification and thereby LMP is required for Matrix-induced IL-1β secretion, LPS-primed BMDCs were pretreated with increasing doses of lysosomal acidification inhibitor BafA1 before Matrix adjuvant treatment. BafA1 blocked the release of IL-1β induced by Matrix-M, Matrix-A, and Matrix-C in a dose-dependent manner (**Fig. 4B**), indicating that lysosomal acidification is crucial for Matrix-induced IL-1β secretion and suggesting a link to LMP. Of note, blockade of lysosomal acidification also prevented the reduction in cell viability observed in response to Matrix adjuvants (**Supplementary Fig. 3A**).

**Figure 4:**
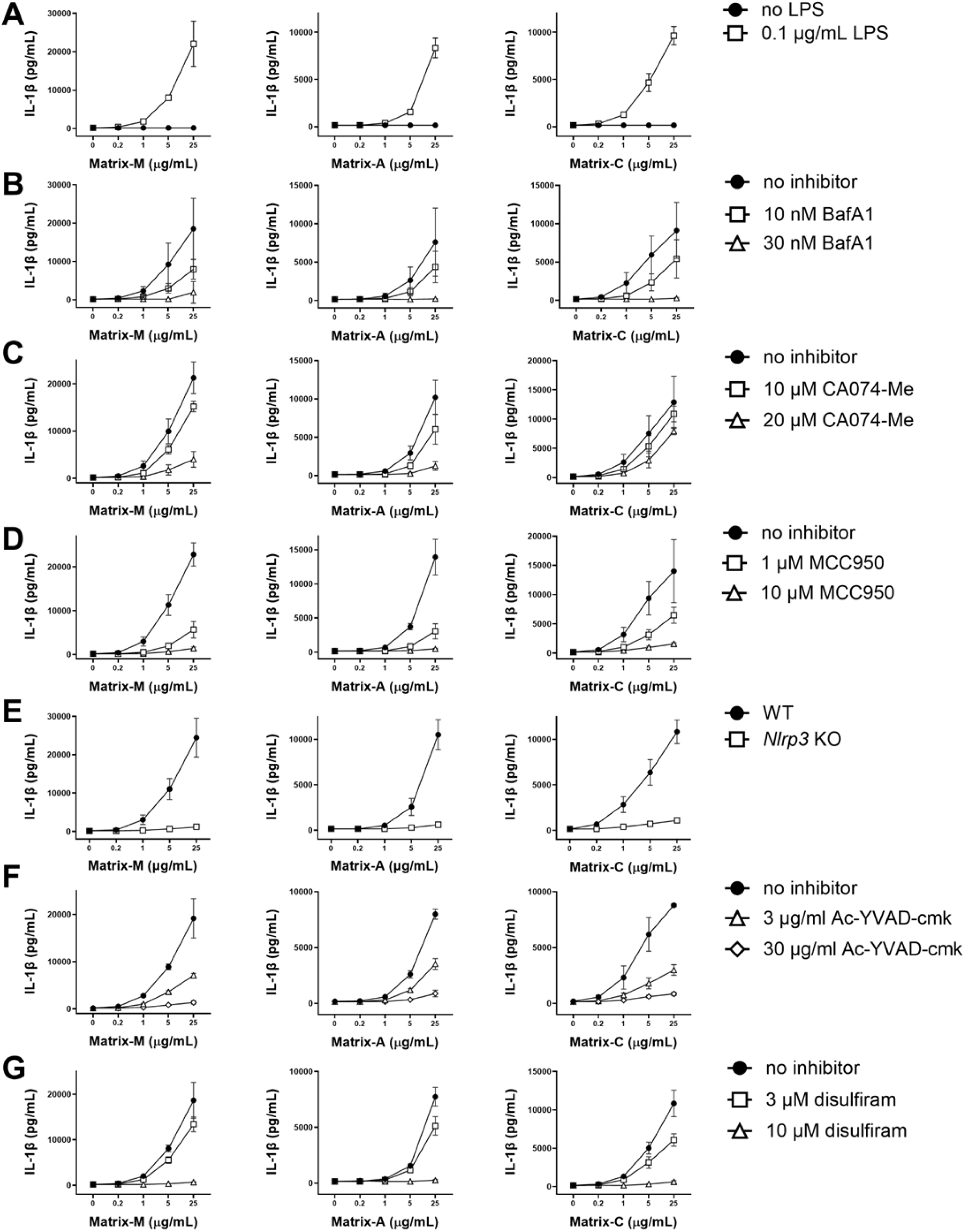
Matrix-M induces NLRP3 inflammasome activation and IL-1β release *in vitro*. BMDCs were either non-primed (**A**) or LPS-primed (**A–G**) and treated with increasing doses of Matrix-M, Matrix-A, or Matrix-C for 5 hours. Released IL-1β levels were measured in the supernatants by ELISA. Before treatment with Matrix adjuvants, cells were pre-treated for 1 h with no inhibitor, V-ATPase inhibitor bafilomycin A1 (BafA1) (**B**), cathepsin B inhibitor CA074-Me (**C**), NLRP3-related inhibitors MCC950 (**D**) and Ac-YVAD-cmk (**F**), or gasdermin D pore formation inhibitor disulfiram (**G**) at the indicated concentrations. Additionally, IL-1β levels were measured in NLRP3-deficient and wild type (WT, C57BL/6) BMDCs after Matrix adjuvant treatment (**E**). Results are shown as the mean ± SD from at least three independent experiments.

One key lysosomal mediator known to have inflammasome activating capability after its release into the cytosol is cathepsin B (CatB)^35^. To test whether CatB contributes to Matrix-induced IL-1β release, LPS-primed BMDCs were pre-treated with the CatB inhibitor CA074-Me before Matrix-M, Matrix-A, or Matrix-C treatment. Inhibition of CatB substantially lowered IL-1β levels induced by Matrix-M and Matrix-A, whereas Matrix-C-induced IL-1β was only marginally affected (**Fig. 4C**).

Among the various inflammasomes, the NLRP3 inflammasome is of particular interest, as lysosomal damage is one of its known activating triggers^36^. To further investigate whether Matrix-induced IL-1β release occurs through NLRP3 inflammasome activation, LPS-primed BMDCs were pretreated with MCC950, an inhibitor of NLRP3 inflammasome assembly. This effectively blocked Matrix-induced IL-1β release (**Fig. 4D**). In line with this, NLRP3-deficient BMDCs failed to release IL-1β upon exposure to Matrix adjuvants (**Fig. 4E**). Given that NLRP3 inflammasome activation results in the activation of caspase-1, which processes pro–IL-1β to mature IL-1β, the dependency of IL-1β release on caspase-1 activation was tested. Pretreatment with the caspase-1 inhibitor Ac-YVAD-cmk resulted in a marked reduction of IL-1β secretion in response to Matrix-M, Matrix-A, and Matrix-C (**Fig. 4F**). This demonstrates that NLRP3 inflammasome activation and caspase-1 are crucial for Matrix-induced IL-1β secretion.

IL-1β lacks the signal sequence required for ER/Golgi-mediated secretion and is consequently released from cells through an unconventional secretion pathway involving gasdermin D (GSDMD) pores^37^. To investigate the potential role of GSDMD in Matrix-induced IL-1β release, LPS-primed BMDCs were pretreated with disulfiram and necrosulfonamide (NSA), both inhibitors of GSDMD that block pore formation by preventing GSDMD oligomerization^38,39^. Both inhibitors dose-dependently blocked IL-1β release induced by Matrix-M, Matrix-A, and Matrix-C (**Fig. 4G, Supplementary Fig. 3B**), showing that Matrix-induced IL-1β is released through GSDMD pores. Inflammasome-mediated GSDMD pore formation can result in either sublytic IL-1β release without causing cell death or lytic pyroptosis^40^. Blocking GDSMD pore formation partially mitigated the reduction in cell viability observed after exposure to Matrix adjuvants (**Supplementary Figure 3C, D**).

### Matrix-M induces IL-18 release independently of NLRP3 inflammasome activation

In addition to IL-1β, IL-18 can be activated through NLRP3 and other inflammasomes. To evaluate the role of Matrix-M, Matrix-A, and Matrix-C in IL-18 release, we measured the levels of IL-18 secreted by non-primed and LPS-primed BMDCs using ELISA. IL-18 release was observed in response to all three adjuvants, with non-primed cells secreting higher levels than LPS-primed cells (**Fig. 5A**). Inhibition of lysosomal acidification again completely blocked Matrix-induced IL-18 release, linking this process to Matrix-induced LMP (**Fig. 5B**). However, and in contrast to observations for IL-1β, Matrix-induced IL-18 secretion was not dependent on NLRP3 inflammasome activation. In fact, inhibition of NLRP3 with the assembly inhibitor MCC950, or the use of NLRP3-deficient BMDCs, resulted in elevated IL-18 secretion responses compared to controls (**Fig. 5C, D**).

**Figure 5:**
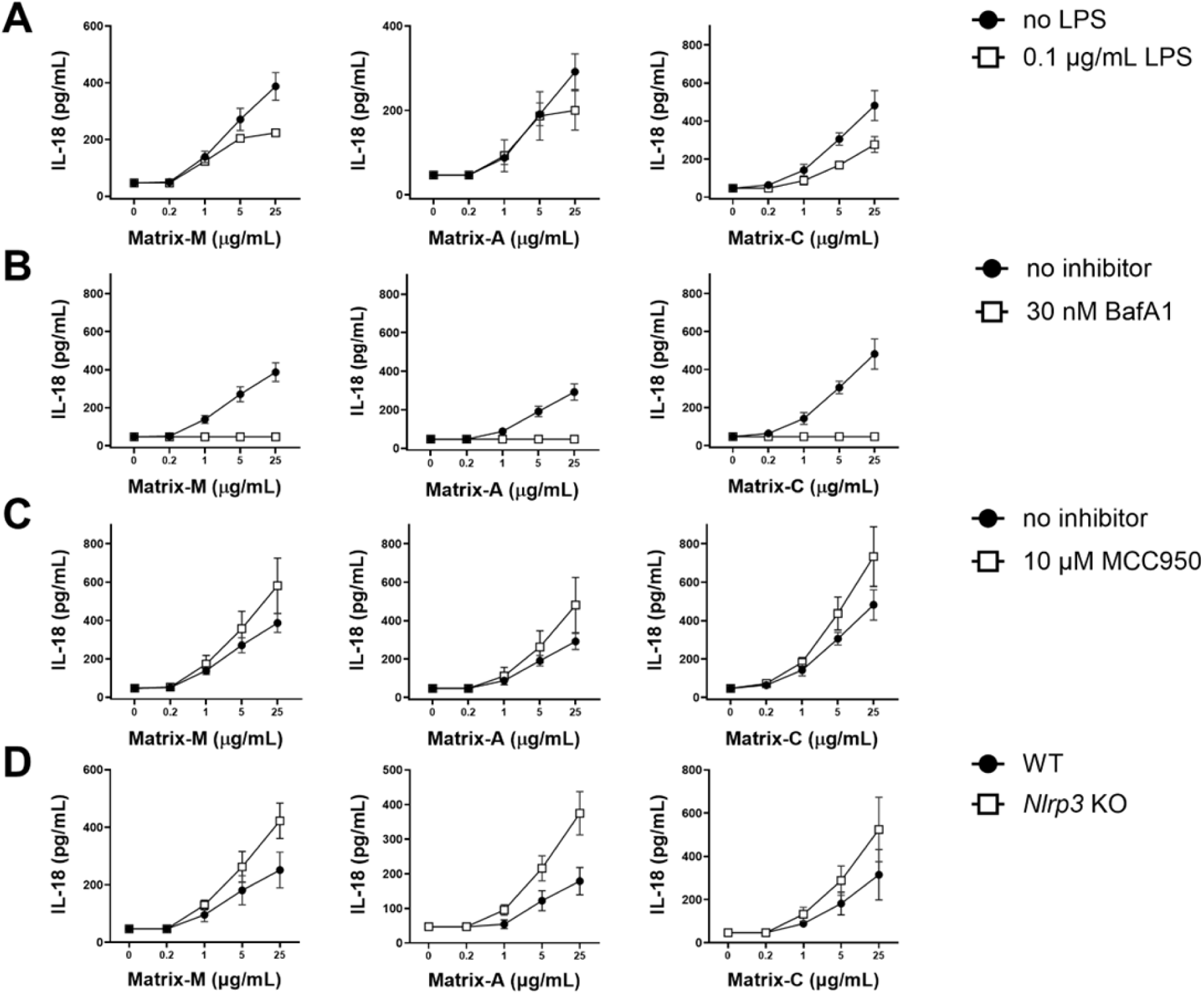
Matrix-M induced IL-18 release is NLRP3 inflammasome independent. BMDCs were either non-primed (**A–D**) or LPS-primed (**A**) BMDCs were treated with increasing doses of Matrix-M, Matrix-A, or Matrix-C for 24 hours. Released IL-18 levels were measured by ELISA. Before treatment with Matrix adjuvants, cells were pre-treated for 1 h with V-ATPase inhibitor bafilomycin A1 (BafA1) (**B**) or NLRP3-assembly inhibitor MCC950 (**C**) at the indicated concentration. Similarly, IL-18 levels were measured in NLRP3-deficient and wild type (WT, C57BL/6) BMDCs after Matrix adjuvant treatment (**D**). Results are shown as mean ± SD of three independent experiments.

### NLRP3 inflammasome activation is not required for Matrix-M adjuvanticity *in vivo*

Given the observed Matrix-induced NLRP3 inflammasome activation *in vitro*, we investigated whether this activation was essential for the adjuvanticity of Matrix-M *in vivo*. Wild type (WT, C57BL/6 mice) and NLRP3-deficient mice (*Nlrp3* KO mice) were immunized in a two-dose immunization regimen with SARS-CoV-2 rS antigen. At days 13 and 20 post-primary immunization as well as at day 28 (7 days post-secondary immunization), similar titers of rS-specific IgG1, IgG2c, and functional human angiotensin converting enzyme (hACE)-2 receptor-blocking antibodies were found in sera of WT and *Nlrp3* KO mice, with both adjuvanted groups showing superior responses compared to their respective unadjuvanted antigen group (**Fig. 6A–C**). Comparable results were observed using Matrix-A and Matrix-C as adjuvants, although *Nlrp3* KO mice displayed higher hACE2 binding inhibition titers than WT mice in the Matrix-A–adjuvanted group at day 13 (**Fig. 6C**).

**Figure 6:**
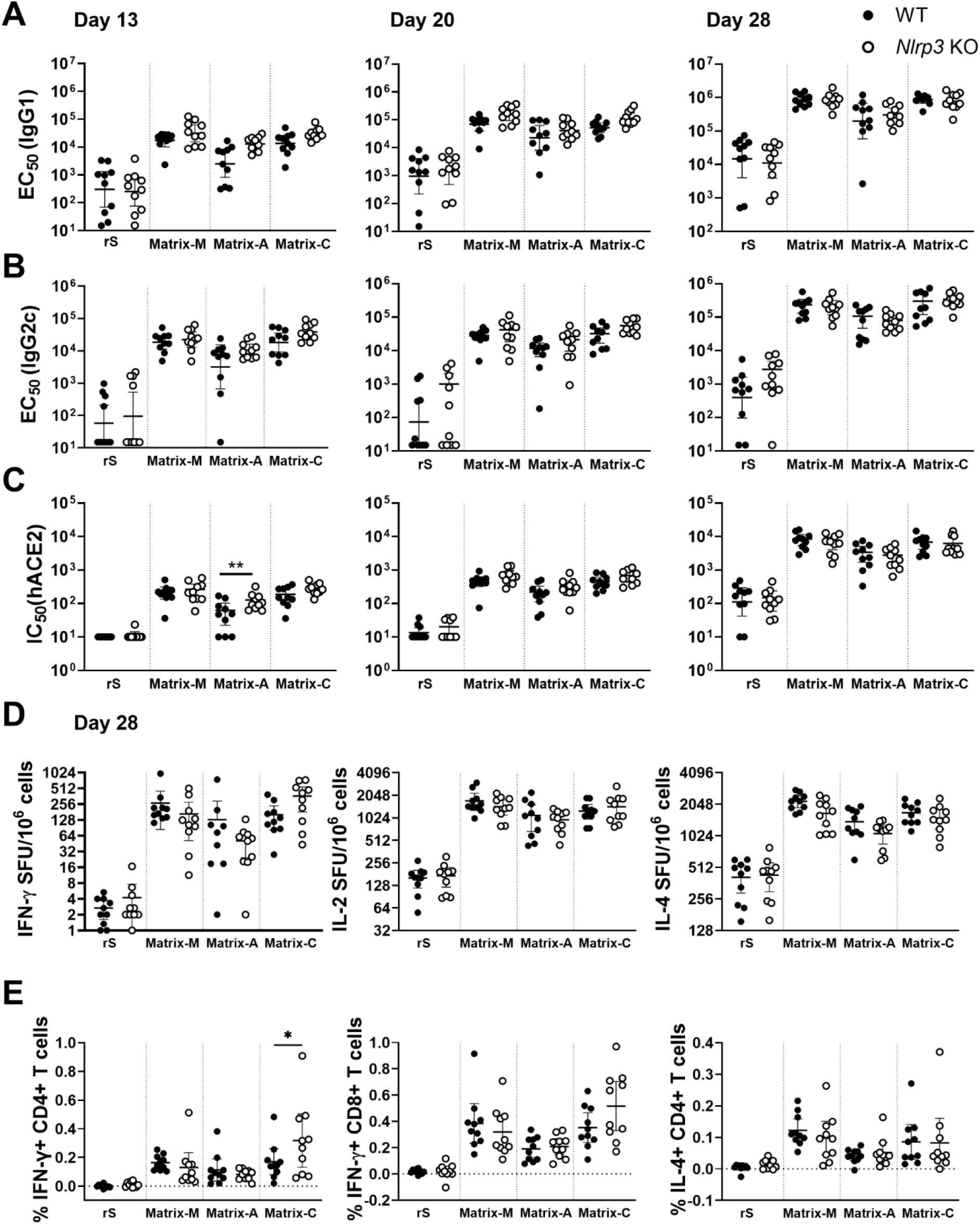
NLRP3 inflammasome activation is not required for Matrix-M adjuvanticity *in vivo*. C57BL/6 (WT) and *Nlrp3* KO mice (n = 10/group) were immunized with 0.1 µg SARS-CoV-2 rS (rS) alone or adjuvanted with 5 μg Matrix-M, Matrix-A, or Matrix-C. Serum samples obtained 13, 20, and 28 days after the primary immunization were evaluated by ELISA for IgG1 (**A**) and IgG2c (**B**) antibody titers against rS protein and for hACE2 receptor-inhibiting antibody titers (**C**). At day 28, splenocytes from individual mice were isolated and the number of cells producing IFN-γ, IL-2, or IL-4 in response to rS protein restimulation was determined by FluoroSpot assay (**D**). Individual titers (ELISA) or spot forming units (SPU)/10^6^ splenocytes are shown as individual symbols, with horizontal bars representing geometric mean values, and error bars represent their 95% confidence intervals. Additionally, splenocytes were analyzed by flow cytometry (intracellular staining) for to access frequencies of rS-specific CD4^+^ and CD8^+^ T cells producing IFN-γ and IL-4 (**E**). Data for each individual mouse response are shown with symbols and group means are represented by horizontal bars. The data (**A–E**) were analyzed for differences between WT and and *Nlrp3* KO mice within each adjuvanted group by one-way ANOVA with Šidák’s multiple comparisons test (*for p < 0.05, ** for p < 0.01).

Analysis of splenocytes producing IFN-γ, IL-2, or IL-4 following rS restimulation at day 28 showed clear adjuvant effects across Matrix-M–, Matrix-A–, and Matrix-C–adjuvanted groups, regardless of the presence or absence of the NLRP3 inflammasome (**Fig. 6D**). ICS analysis revealed similar frequencies of antigen-specific IFN-γ–producing CD4^+^ and CD8^+^ T cells, as well as IL-4–producing CD4^+^ T cells in WT and *Nlrp3* KO mice in the non-adjuvanted and Matrix-M– and Matrix-A–adjuvanted groups (**Fig. 6E**). In the Matrix-C–adjuvanted groups, a higher frequency of antigen-specific IFN-γ–producing CD4^+^ T cells was observed in *Nlrp3* KO mice, while similar frequencies of IFN-γ–producing CD8^+^ and IL-4–producing CD4^+^ T cells were found in Matrix-C–adjuvanted WT and *Nlrp3* KO mice. Collectively, the data demonstrate that Matrix-M, Matrix-A, and Matrix-C show potent adjuvant effects *in vivo* both in the presence and absence of the NLRP3 inflammasome.

### Matrix-M enables antigen cross-presentation in BMDCs

The localization of Matrix-M inside lysosomes together with its observed pore-forming activity, could allow exogenous antigens to reach the cytosol. This, in turn, may enable endocytosed antigens to enter the MHCI processing pathway, thereby promoting antigen cross-presentation, a mechanism that has been exemplified by other SBAs^15,24,27^. Given that Matrix-A and Matrix-C particles in Matrix-M are not physically linked to antigens and thus Matrix-M does not serve as a direct antigen delivery system, we first evaluated the colocalization of the antigen with lysosomes and Matrix-M as a prerequisite condition for cross-presentation. LysoTracker-loaded BMDCs were incubated with BODIPY-conjugated Matrix-M together with Alexa Fluor 647–labeled SARS-CoV-2 rS antigen and monitored by time-lapse spinning-disk confocal live-cell imaging. PCC analysis revealed a marked time-dependent increase in the degree of colocalization between LysoTracker and the antigen, confirming antigen internalization into the lysosomes (**Fig. 7A**). In parallel, colocalization between Matrix-M and the antigen was also confirmed (**Fig. 7B**). These results demonstrate that both Matrix-M and the antigen accumulate within lysosomes upon cellular uptake.

**Figure 7:**
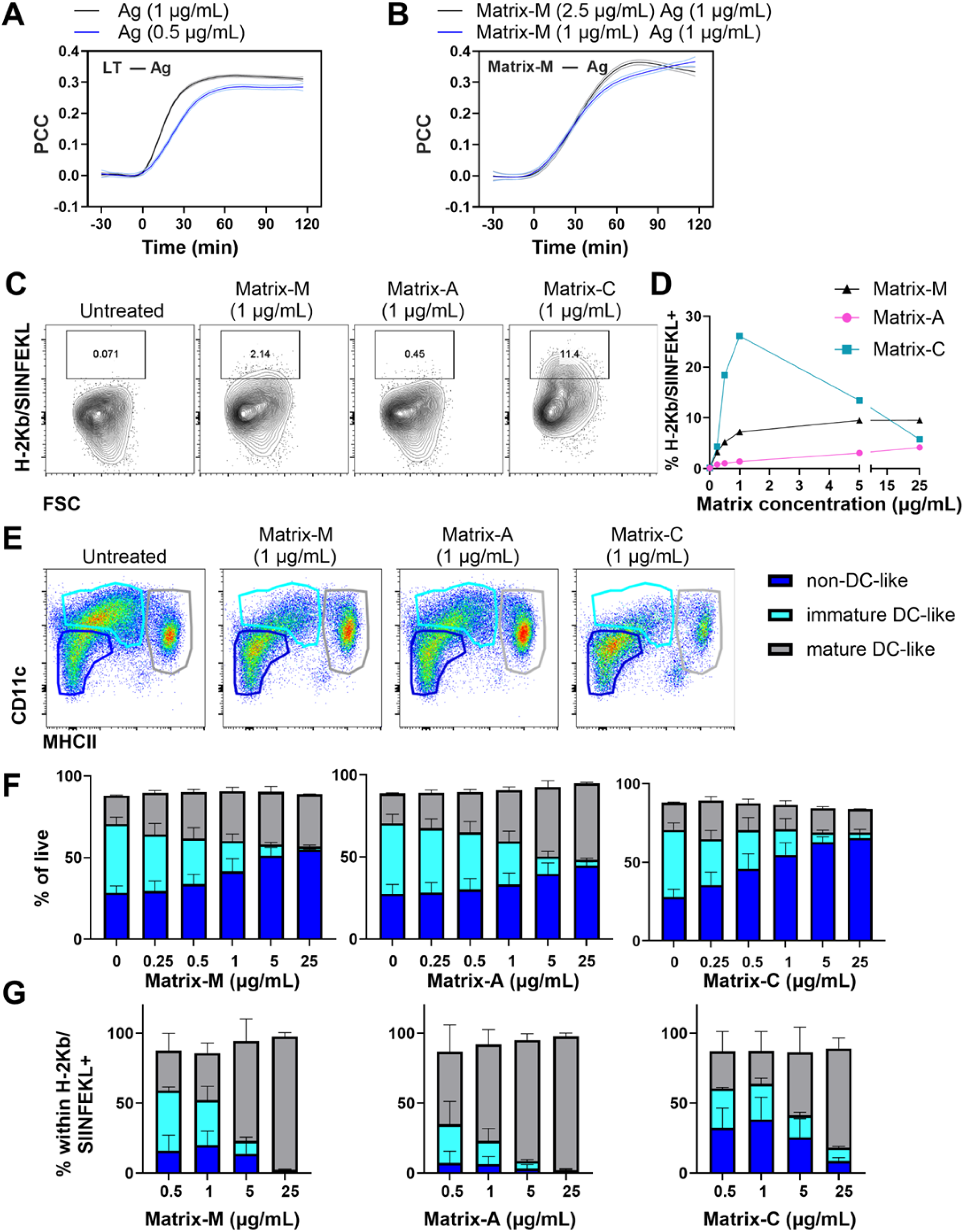
Matrix-M enables antigen cross-presentation in BMDCs. BMDCs were loaded with LysoTracker Red DND-99, washed and imaged using spinning-disk confocal microscopy for 30 minutes. Following the initial pre-bleaching image acquisition, cells were treated with Alexa Fluor 647–labeled rS (0.5 or 1 μg/mL) and BODIPY-Matrix-M (1 or 2.5 μg/mL) and live-cell imaging continued for an additional 120 min. Pearson’s correlation coefficients (PCC) for the channel combination LysoTracker and Alexa Fluor 647 **(A)**, and BODIPY and Alexa Fluor 647 **(B)** were determined as indicators of lysosome–antigen or Matrix‒M–antigen colocalization, respectively. Data in panels **A** and **B** are presented as smoothed averages ± SEM, pooled from at least three experiments, each performed in triplicates. **(C–G)** BMDCs were exposed to 0.8 mg/mL OVA and indicated concentrations of Matrix-M, Matrix-A, or Matrix-C for 24 hours. Cells were analyzed by flow cytometry for surface markers and for H-2kB bound to SIINFEKL complexes, with the latter indicative of antigen cross-presentation. The percentage of viable BMDCs positive for H-2kB/SIINFEKL complexes is shown as representative contour plots **(C)** and corresponding graphs **(D)** for untreated and Matrix-treated groups. **(E)** Representative flow cytometry plots and **(F)** graphs show the proportion of major subpopulations within BMDC culture, identified as non-DC-like cells (CD11c^−^ MHCII^−^, dark blue), immature DC-like cells (CD11c^+^ MHCII^−/lo^, turquoise) and mature DC-like cells (CD11c^+^ MHCII^hi^, gray). **(G)** The percentage of various BMDC subpopulations within cells positive for H-2Kb/SIINFEKL complexes are shown. Data in panels **C–E** are representative of three independent experiments, while graphs in **F** and **G** show pooled data from three independent experiments, presented as the mean ± SD.

The ability of Matrix-M to induce antigen cross-presentation was subsequently investigated using the 25-D1.16 monoclonal antibody, which specifically recognizes cell surface complexes of the OVA peptide SIINFEKL bound to the MHCI molecule H-2Kb. BMDCs treated with OVA and Matrix-M for 24 h showed a Matrix-M–induced cross-presentation (**Fig. 7C, D**). Similarly, antigen cross-presentation was observed in BMDCs after treatment with Matrix-A and Matrix-C (**Fig. 7C, D**). When comparing the cross-presentation–inducing potential of the Matrix adjuvants, Matrix-C elicited the highest frequency of antigen cross-presenting cells at Matrix concentrations of 5 µg/mL or lower, followed by Matrix-M, while Matrix-A induced the lowest level of cross-presentation. Both Matrix-M and Matrix-A promoted a concentration-dependent increase in the frequency of antigen cross-presenting cells, whereas treatment with Matrix-C at concentrations above 5 µg/mL resulted in a reduced frequency of cross-presenting BMDCs. Phenotypic analysis of BMDCs, categorized into non–dendritic cell (DC)-like, immature DC-like, and mature DC-like populations (gating strategy shown in **Supplementary Figure 4**), showed a decrease in the proportion of immature DC-like cells after Matrix treatment (**Fig. 7E, F**). Antigen cross-presentation was primarily driven by mature DC-like cells at 5 and 25 µg/mL Matrix concentrations, irrespective of the Matrix formulation (Matrix-M, Matrix-A, or Matrix-C) used (**Fig. 7G**). Furthermore, the effect of Matrix adjuvants on BMDC activation was investigated, with Matrix-M and Matrix-A showing the strongest effect, as indicated by increased frequencies of MHCII^+^ and CD86^+^ cells (**Supplementary Fig. 5A–H**). At higher concentrations (5 and 25 µg/mL), Matrix-A treatment led to higher proportions of MHCII^+^ and CD86^+^ BMDCs compared to Matrix-M (**Supplementary Fig. 5C, G**). In line with the *in vitro* finding that the Matrix-induced IL-1β secretion depends on NLRP3 inflammasome activation, the upregulation of activation markers MHCII and CD86 on BMDCs was reduced in NLRP3-deficient cells following Matrix-M, Matrix-A, or Matrix-C treatment (**Supplementary Figure 6A, B**). Altogether, these results suggest that the combination of Matrix-C and Matrix-A in Matrix-M effectively merges Matrix-C’s potential for antigen cross-presentation with Matrix-A’s ability to promote APC activation.

### Lysosomal acidification and proteasomal degradation contribute to Matrix-induced antigen cross-presentation

To assess the ability of APCs to cross-prime CD8^+^ T cells, BMDCs were co-cultured with B3Z cells, a co-stimulation independent reporter T-cell line^41^. The B3Z cross-presentation assay, in which the presentation of SIINFEKL via H-2Kb activates T-cell receptor signaling inducing β-galactosidase expression, showed a concentration dependent induction of cross-priming, with a higher potential of Matrix-C compared to Matrix-M and Matrix-A (**Fig. 8A**). Importantly, the exposure of cells to Matrix-M, Matrix-A or Matrix-C did not alter MHCI presentation or cell viability, as presentation of pulsed SIINFEKL was similar or slightly reduced within the tested concentration range (**Fig. 8B**). Blockade of lysosomal acidification showed that Matrix-induced cross-presentation was dependent on a low lysosomal pH, suggesting a link to LMP (**Fig. 8C, D**), consistent with the observed inhibition of dextran release by BafA1 (**Fig. 3B**). For the cross-presentation of exogenous antigens, two main intracellular pathways, the “endosomal” and “cytosolic,” have been proposed^42^. Given that Matrix-induced dextran release indicated antigen translocation into the cytosol, the role of proteasomal degradation, a key component of the cytosolic pathway in cross-presentation, was examined. Inhibition of proteasomal activity by epoxomicin resulted in a strong reduction in cross-presentation induced by Matrix-M, Matrix-A, and Matrix-C, confirming that Matrix-induced antigen translocation into the cytosol underlies Matrix-driven cross-presentation (**Fig. 8E, F**).

**Figure 8:**
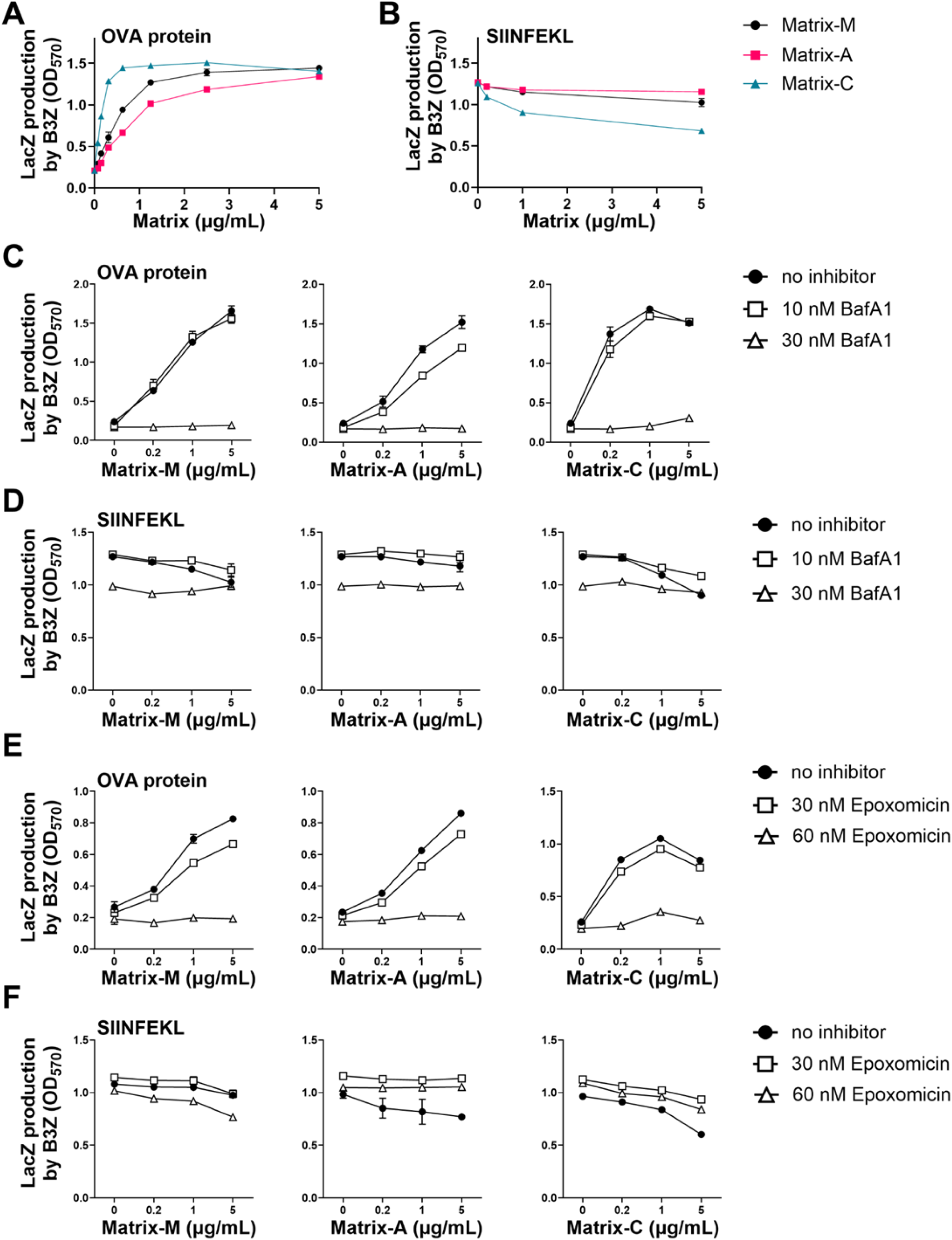
Lysosomal acidification and proteasomal degradation contribute to Matrix-induced antigen cross-presentation. BMDCs were exposed to 80 µg/mL OVA and indicated concentrations of Matrix-M, Matrix-A, or Matrix-C for 5 h (**A, C, E**). In control experiments, 5 ng/mL SIINFEKL instead of OVA was added (**B, D, F**). Cells were co-cultured for 18 h with B3Z cells, which produce β-galactosidase upon TCR recognition of SIINFEKL presented on H-2kB. BMDCs were pretreated for 1 h with bafilomycin A1 (BafA1) to inhibit lysosomal acidification (**C, D**) or with epoxomicin (**E, F**) to inhibit proteasomal activity. Results shown are representative of two independent experiments, with data presented as mean values of triplicates ± SD.

### Matrix-M induces dose-dependent CD8^+^ T-cell responses *in vivo*

To further evaluate the ability of Matrix-M to induce antigen cross-presentation and generate antigen-specific CD8^+^ T-cell responses, two different approaches were employed. First, antigen cross-presentation was assessed *ex vivo* by co-culturing sorted CD11c^+^ DCs, which were loaded *in vivo* with OVA either in presence or absence of Matrix-M, with naïve OT-I CD8^+^ T cells that carry an OVA peptide-specific T-cell receptor. Immunization with OVA adjuvanted with Matrix-M resulted in enhanced proliferation and activation of naïve OVA-specific CD8^+^ T cells in *ex vivo* co-cultures (**Fig. 9A–D**). Analysis of the CD11c-sorted DCs showed a shift in proportions of DC subsets, with an increase in monocyte-derived DCs (moDCs) and a decrease in conventional DC1 and DC2 (cDC1 and cDC2) subpopulations in the group receiving Matrix-M (**Supplementary Fig. 7A, B**). Moreover, Matrix-M treatment led to increased activation of DCs in the draining lymph nodes (dLNs) (**Supplementary Fig. 7C**).

**Figure 9:**
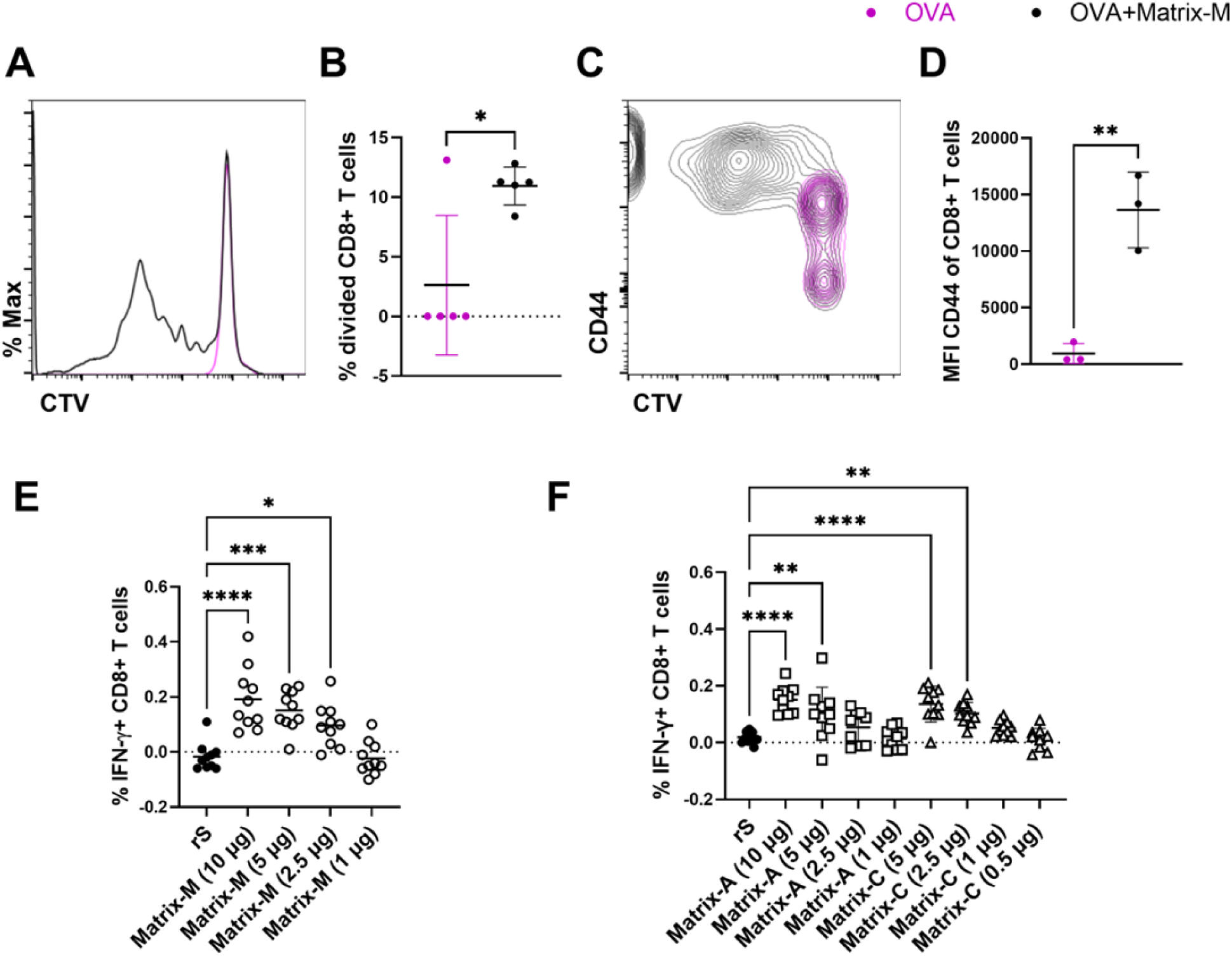
Matrix-M generates dose dependent CD8^+^ T-cell responses *in vivo*. C57BL/6 mice were immunized with OVA alone or adjuvanted with 5 µg Matrix-M. Pools of DCs isolated from the dLNs were co-cultured for 3 days with naïve CD8^+^ OT-I T cells. Representative histograms for CellTrace Violet (CTV) dilution are shown (**A**); these were used to calculate the percentage of divided cells by FlowJo proliferation analysis (**B**). Surface expression (median fluorescence intensity; MFI) of the T-cell activation marker CD44 within CD8^+^ T cells was analyzed, as displayed in the representative contour plot and corresponding graph (**C, D**). Each symbol represents one individual pool, with the solid horizontal bars representing group means ±SD. Unpaired t-test was applied for comparisons between the two groups (**B, D**). (**E**, **F**) BALB/c mice (n = 10/group) were immunized with 0.1 µg SARS-CoV-2-rS alone or adjuvanted with the indicated doses of Matrix-M (**E**), Matrix-A, or Matrix-C (**F**). Splenocytes from individual mice were isolated to determine the frequency of CD8^+^ T cells that produce IFN-γ after rS protein stimulation. Each symbol represents an individual mouse; horizontal bars represent group means. One-way ANOVA with Tukey’s multiple comparisons test was applied to compare the groups to antigen only (p-values * for p <0.05, ** p <0.01, *** p<0.001, **** p<0.0001).

In a second approach, the effect of Matrix-M adjuvant on CD8^+^ T-cell responses was investigated *in vivo* using a two-dose immunization scheme with SARS-CoV-2 rS protein, where different doses of Matrix-M were administered. ICS for IFN-γ demonstrated a dose dependent induction of rS-specific CD8^+^ T-cell responses (**Fig. 9E**). In addition, a dose response was observed for the generation of specific serum antibodies, cellular responses, and CD4^+^ T-cell responses (**Supplementary Fig. 8A–D**). Similarly, a dose-dependent induction of antigen-specific CD8^+^ T cells was observed when varying doses of Matrix-A and Matrix-C adjuvants were used for rS immunizations (**Fig. 9F**). Although non-adjuvanted rS immunization did not elicit detectable rS-specific CD8^+^ T cells, the addition of 2.5 µg Matrix-M and Matrix-C or 5 µg Matrix-A was sufficient to induce CD8^+^ T-cell responses. Altogether these data demonstrate the ability of Matrix-M, as well as that of Matrix-A and Matrix-C, to induce specific CD8^+^ T cells responses to exogenous antigens, as demonstrated with two distinct antigens, OVA and SARS-CoV-2 rS protein.

## Discussion

Matrix-M was here studied alongside its constituent subcomponents, Matrix-A and Matrix-C particles, to investigate intracellular events following cellular uptake and their impact on the adjuvant effect of Matrix-M. Our findings demonstrate that Matrix-M, Matrix-A, and Matrix-C all can induce LMP in BMDCs *in vitro*, a process that strongly depends on an acidic lysosomal environment. Further investigations revealed that Matrix-induced LMP triggered NLRP3 inflammasome activation, which was essential for IL-1β secretion by BMDCs, but was not required for Matrix-M’s adjuvanticity *in vivo*. Furthermore, Matrix-induced LMP enabled antigen cross-presentation leading to the induction of antigen-specific CD8^+^ T-cell responses.

Matrix-M adjuvant is a key component in licensed vaccines for COVID-19 and malaria and in other vaccine candidates due to its potent adjuvant properties combined with a favorable safety profile^1–6,11^. Matrix-M is composed of 85% Matrix-A and 15% Matrix-C particles (based on saponin weight), both derived from distinct *Quillaja* saponin fractions^43^. *Quillaja* saponins share a common quillaic acid triterpene aglycone chemical structure, with variations in glycoside side chains.

LMP can cause the release of lysosomal contents, such as various proteases, into the cytosol, triggering programmed cell death or inflammatory responses^44^. SBAs such as QS-21 and ISCOMATRIX have been shown to induce LMP, potentially enhancing adjuvant activity by promoting innate immune activation (for example, through Syk kinase or inflammasome activation) and facilitating antigen cross-presentation^14–16,21^. Here, we demonstrate that Matrix-M similarly induces LMP in BMDCs. Time-lapse spinning-disk confocal imaging revealed that Matrix-C has a stronger LMP-inducing potential than Matrix-A and, to some extent, Matrix-M. In line with this finding, Matrix-A showed the highest colocalization with lysosomes in BMDCs, most likely due to a lower degree of LMP. Our previous findings showed that Matrix-A particles (and presumably also Matix-C, although not specifically studied to date) rapidly disassemble both at the injection site and in the draining lymph node, separating into saponins and cholesterol^13^. It is known that saponins, when not formulated together with cholesterol and phospholipids, have lytic activity^11^. Specifically, the long fatty acyl chain in Fraction-C saponins is thought to integrate into the plasma membranes, contributing to their destabilization.

The proposed disassembly of Matrix-A and Matrix-C particles in lysosomes is likely driven by the acidic environment and/or the enzymatic activity within these organelles. Of note, cytosolic leakage of the 3-kDa dextran particles by Matrix-C was only partially blocked by inhibition of the V-ATPase proton pump, which is responsible for maintaining low lysosomal pH. This suggests that Matrix-C retains some LMP-inducing capacity even under non-acidic conditions. This potential for inducing LMP under non-acidic conditions may explain why Welsby *et al.*, despite observing similar cytosolic translocation of 3-, 10-, and 40-kDa dextran particles after QS-21 exposure, could not inhibit the lysosomal membrane leakage using the V-ATPase proton pump inhibitor BafA1^16^. The potential degradation mechanism of Matrix particles and the structure of the resulting disassembled saponins, cholesterol, phospholipids, and associated fragments were beyond the scope of this study but remain areas of interest for future research. Additionally, blocking lysosomal acidification inhibited inflammasome activation and antigen cross-presentation, connecting these processes to Matrix-induced LMP.

NLRP3 inflammasome activation mediated IL-1β release in Matrix-treated BMDCs, which is in line with observations for other SBAs such as QS-21 or ISCOMATRIX^21,22,45,46^. However, without priming by LPS or MPLA, there was no IL-1β release, strongly suggesting that Matrix-M, similar to other SBAs, primarily functions as the second signal in NLRP3 inflammasome activation. As expected, Matrix-induced release of IL-1β is mediated through GSDMD pores^47^. However, Matrix-induced cell death is not altered when GSDMD pore formation is inhibited, indicating that IL-1β release occurs independently of conventional pyroptosis and follows a distinct pathway from cell death^48,49^.

Matrix-induced IL-18 release further highlights the significance of lysosomal acidification and LMP. Nonetheless, the priming and NLRP3 inflammasome activation steps necessary for IL-1β release were found to be non-essential for IL-18 release and may even inhibit it. The lack of requirement for LPS priming in Matrix-induced IL-18 secretion confirms a constitutive expression of pro-IL-18 by BMDCs and points to a possible competition between TLR4-induced, Matrix-induced IL-1β and IL-18 pathways. These intersecting pathways may involve shared proteases, such as caspase-1, needed for pro-GSDMD cleavage into its pore-forming form^47^, or caspase-8, which could activate caspase-1 upstream of NLRP3 or directly cleave pro–IL-1β/IL-18^50–52^. Other studies using BMDCs and murine or human monocyte-derived macrophages indicate that IL-18 release generally requires priming and NLRP3 activation^21,22^. Notably, comparing findings across different cell types can be challenging. In animal studies, IL-18 has been shown to play a key role for the adjuvanticity of SBAs by driving an early IFN-γ response critical for the generation of Th1-biased CD4^+^ T-cell responses (AS01), CD8^+^ T cell responses (ISCOMATRIX) and Th1 isotype switching, leading to antigen-specific IgG2c antibody responses (AS01, ISCOMATRIX)^22,23^. The ability of Matrix-M to induce IL-18 secretion in BMDCs (in an LPS/NLRP3-independent manner), along with its Th1-skewed immune profile, suggest that IL-18 might similarly contribute to the mechanism of action of Matrix-M.

Despite the role of the NLRP3 inflammasome in Matrix-induced IL-1β release *in vitro*, no major differences were observed in the humoral or cellular responses between WT and *Nlrp3* KO mice following SARS-CoV-2 rS vaccination with Matrix-M, Matrix-A, or Matrix-C. These results are consistent with findings for ISCOMATRIX, where NLRP3-deficient mice showed normal IgG1 and IgG2c antibody titers and antigen-specific CD8^+^ T-cell responses to OVA vaccination^22^. Alternative inflammasome receptors may compensate for the absence of NLRP3 *in vivo*, as deletion of other inflammasome components, such as ASC and caspase-1/11, resulted in a partial reduction of immune responses in the aforementioned ISCOMATRIX experiments. By contrast, elevated Th1 and Th2 responses and antigen-specific IgG1 and IgG2c antibody titers were found in the absence of NLRP3 for the adjuvant effect of QS-21^21^. Altogether, these data indicate that the adjuvant effect of Matrix-M *in vivo* is robustly observed, both in the presence and absence of the NLRP3 inflammasome.

While low doses of all three Matrix adjuvants induce LMP and NLRP3 inflammasome activation, higher doses may cause excessive LMP that could downregulate NLRP3 activation and ultimately lead to cell death^53,54^. Matrix-C, in particular, triggers marked LMP followed rapidly by cell death at relatively low concentrations, whereas Matrix-A requires considerably higher doses to achieve similar effects. This difference could be due to induction of a more efficient plasma membrane damage repair response or reduced membrane damage by Matrix-A. Cell death-dependent release of DAMPs is considered part of SBA mechanism of action^55^.

Matrix-M injection has been shown to activate DCs in the dLNs of mice, as evidenced by elevated expression of the co-stimulatory molecule CD86^12,56^. Interestingly, while Matrix-C showed the greatest potential for LMP, we observed little activation of APCs by Matrix-C in our *in vitro* model. By contrast, Matrix-M and, more notably, Matrix-A were found to induce a robust activation of BMDCs, as shown by elevated expression of MHCII and CD86. Interestingly, findings from other SBAs are somewhat conflicting, with QS-21 having been demonstrated to upregulate expression of CD86 and HLA-DR in human moDCs^16^, while Matrix-C and ISCOMATRIX both failed to do so in BMDCs^24,57^. One possible explanation for the observed activation of Matrix-M and Matrix-A treated BMDCs could be that Matrix-M and Matrix-A do not cause excessive LMP, allowing cells to bypass pyroptosis and enter a hyperactivated state following inflammasome activation^40^. These findings suggest that the effectiveness of Matrix-M likely stems from its ability to combine strong cross-presentation capacity (largely driven by Matrix-C) and robust CD4^+^ T-cell activation (mainly driven by Matrix-A).

Of note, GM-CSF-differentiated BMDCs are a heterogeneous population, comprising conventional DCs and monocyte-derived macrophages^58^. Treatment of BMDCs with Matrix-M, Matrix-A, or Matrix-C led to a concentration-dependent reduction in the frequency of immature DCs, accompanied by an increase in both non–DC-like cells and mature DC-like cells for Matrix-M and Matrix-A and by an increase in non–DC-like cells for Matrix-C. This shift could be due to a preferential cell death of immature DCs, a differentiation of immature DCs into mature DCs, or a proliferation of the increasing populations. Notably, within the BMDC population driving antigen cross-presentation, mature DC-like cells were the most prominent subset. These results differ from previous works by den Brok *et al.* and Huis in ‘t Veld *et al.*, which identified the MHCII^lo^ CD11b^hi^ DC subset, corresponding to immature DC-like cells in this study, as the main antigen cross-presenting population^24,26^. However, in those studies, the BMDCs were sorted before Matrix-C exposure, whereas our data show that this population of immature DC-like cells diminishes after addition of Matrix-C, indicating a possible differentiation into mature DC-like cells in aforementioned studies.

The ability of an adjuvant to promote antigen cross-presentation, for example by enabling translocation of the antigen from the lysosome to the cytosol, is essential for generating antigen-specific CD8^+^ T-cell responses in response to protein-based vaccines. We found that Matrix-M, as well as Matrix-A and Matrix-C, induced antigen cross-presentation as a direct downstream effect of LMP in our *in vitro* BMDC model. Matrix-C showed the highest potential, followed by Matrix-M and Matrix-A. These findings are in line with previously reported *in vitro* antigen cross-presentation induction by QS-21, ISCOMATRIX, and Matrix-C^14,15,24–27^. Cross-presentation potential has also been attributed to the immunoactive saponin fractions such as Supersap, Vaxsap and Fraction-C, but not to crude saponin fractions^24^. Matrix-A and Matrix-C both contain distinct immunoactive saponin fractions, and this study is, to our knowledge, the first to demonstrate the cross-presentation–inducing capacity of Matrix-A, the constituent saponins of which lack a fatty acyl chain. The proteasome dependency of Matrix-induced antigen cross-presentation confirmed the utilization of the cytosolic pathway by Matrix-M, Matrix-A, and Matrix-C, consistent with previous findings for ISCOMATRIX and Matrix-C^15,24,27^.

Our study demonstrated that DCs sorted from the dLNs after immunization with OVA and Matrix-M could activate OT-I CD8^+^ T cells *ex vivo*. In the Matrix-M–adjuvanted group, moDCs, absent in the dLNs of the antigen-only immunized mice, accounted for approximately one-third of the CD11c^+^/MHCII^+^ DC population. This is in line with data generated using the AS01 adjuvant system, showing that CD11c^+^ DCs are essential for the generation of CD8^+^ T-cell responses *in vivo*^59,60^. Similarly, CD11c^+^ cells, but not CD11c-cells, isolated from the dLNs of mice co-administered with OVA and Matrix-C, were shown to activate OVA peptide-specific OT-I CD8^+^ T cells *ex vivo*^24^.

In alignment with our *in vitro* and *ex vivo* findings, SARS-CoV-2 rS vaccines adjuvanted with either Matrix-M, Matrix-A, or Matrix-C elicited rS-specific CD8^+^ T-cell responses *in vivo* in a dose-dependent manner. Non-adjuvanted rS vaccination failed to induce rS-specific CD8^+^ T-cell responses. A minimum dose of 2.5 µg Matrix-M or Matrix-C, or 5 µg of Matrix-A was required to induce detectable antigen-specific CD8^+^ T-cell responses. This further indicates synergistic effects between Matrix-A and Matrix-C in Matrix-M. Previous investigations have demonstrated that Matrix-C and Matrix-M can generate CD8^+^T-cell responses in mice^24,28,29^. Furthermore, Matrix-A was capable of inducing antigen-specific CD8^+^ T-cell responses *in vivo*.IThese observations invite future research to explore how acyl chain length and other structural properties of saponins in Matrix formulations modulate the extent of LMP and downstream effects, including antigen cross-presentation.

Thus, Matrix-M clearly has the potential to induce antigen cross-presentation and CD8^+^ T-cell responses in pre-clinical models. In clinical studies, the NVX-CoV2373 vaccine, containing SARS-CoV-2 rS protein and Matrix-M, has demonstrated the ability to trigger rS-specific CD8^+^ T-cell responses in 10-50% of vaccine recipients^10,61^. Notably, the frequencies of rS-specific memory CD8^+^ T cells were comparable to those observed in individuals who had recovered from SARS-CoV-2 infection, at the 6-month follow up. An explanation for the lower frequency of CD8^+^ T-cell responses in the human versus the mouse data could be the relatively higher Matrix-M dose administered to mice in relation to the body weight. In addition to vaccine-induced anti-viral responses, CD8^+^ T cells play a crucial role in therapeutic cancer vaccines. Matrix-M may, therefore, also be applicable in areas outside infectious disease vaccines, as tested for other SBAs in clinical trials (NCT05638698)^62^.

Here, we demonstrate that Matrix-M and co-administered antigens enter lysosomes together, where Matrix-M then induces LMP in an acidification-dependent process, thus enabling antigen translocation into the cytosol, NLRP3 inflammasome activation, antigen cross-presentation, and the generation of antigen-specific CD8^+^ T-cell responses. In addition, even in the event of LMP, Matrix-M induces high antibody titers and antigen-specific CD4^+^ T-cell responses, indicating a balanced antigen presentation via both MHCI and MHCII pathways. The combination of Matrix-C and Matrix-A in Matrix-M thus confers a dual advantage: the potent ability of Matrix-C to enable antigen cross-presentation and the capacity of Matrix-A to strongly activate APCs. These distinct characteristics likely explain the high overall adjuvant activity of Matrix-M and motivate further evaluation for targeted applications.

## Acknowledgements

We thank Louis Fries for insightful discussions and valuable comments on the manuscript. We thank the staff at SVA for their expert technical help. MES acknowledges support from the SciLifeLab Fellows program and the Swedish Foundation for Strategic Research (FFL18-0165). The funder played no role in study design, analysis, or the writing of this manuscript.

## Author contributions

BZ, BC, LS, and CLA conceptualized the project. BZ, BC, and CLA designed and performed experiments. JE, ES, PÖ, EB, ILO, and AEP performed experiments. EA, LL, JA, NH, PH, and JB developed critical reagents. BZ, BC, JE, KLB, MES, AEP, LS, and CLA analyzed and primarily discussed data. BZ, BC, and CLA wrote the initial manuscript before review by JE, MES, and AEP. All authors read and approved the manuscript.

## Data availability

All data generated or analyzed during this study are included in this published article and its supplementary information files, with the exception when representative data or summarized data from repeated experiments are shown. Raw data are available upon reasonable request to the corresponding author.

