## Supplementary Figures for "Matrix-M® adjuvant triggers inflammasome activation and enables antigen cross-presentation through induction of lysosomal membrane permeabilization"

### Supplementary Figure 1

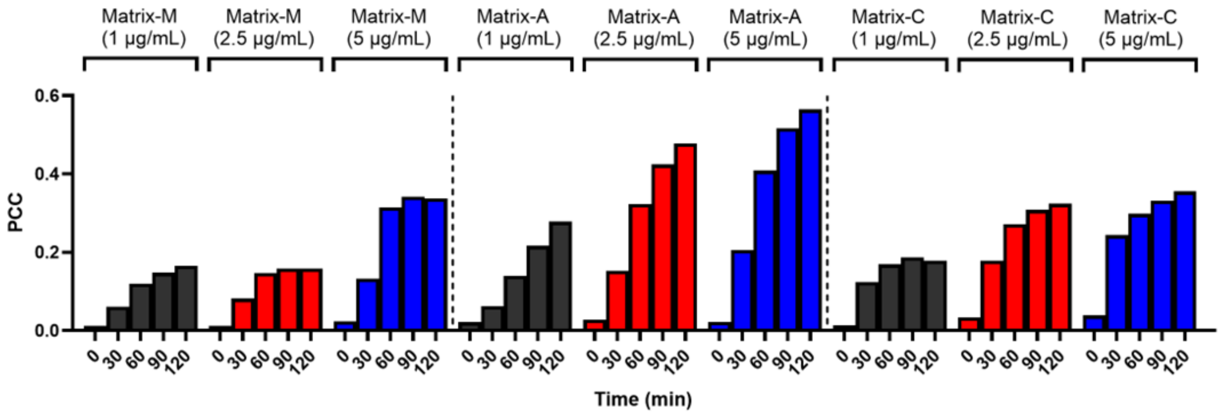


**Supplementary Figure 1:** Matrix adjuvant-lysosome colocalization analysis

Spinning-disk confocal microscopy (live-cell imaging) was applied to study the localization of Matrix adjuvant in lysosomes. LysoTracker-loaded BMDCs were incubated with BODIPY-Matrix-M, Matrix-A, or Matrix-C and imaged for 120 min every 3 minutes. Pearson's correlation coefficient (PCC) for the channel combination BODIPY and LysoTracker was calculated as a measure for the colocalization of Matrix adjuvants with lysosomes. Graphs show the PCC for the respective Matrix adjuvant type and concentration at selected time points. Mean values from at least three replicates are shown.

**Supplementary Figure 2**

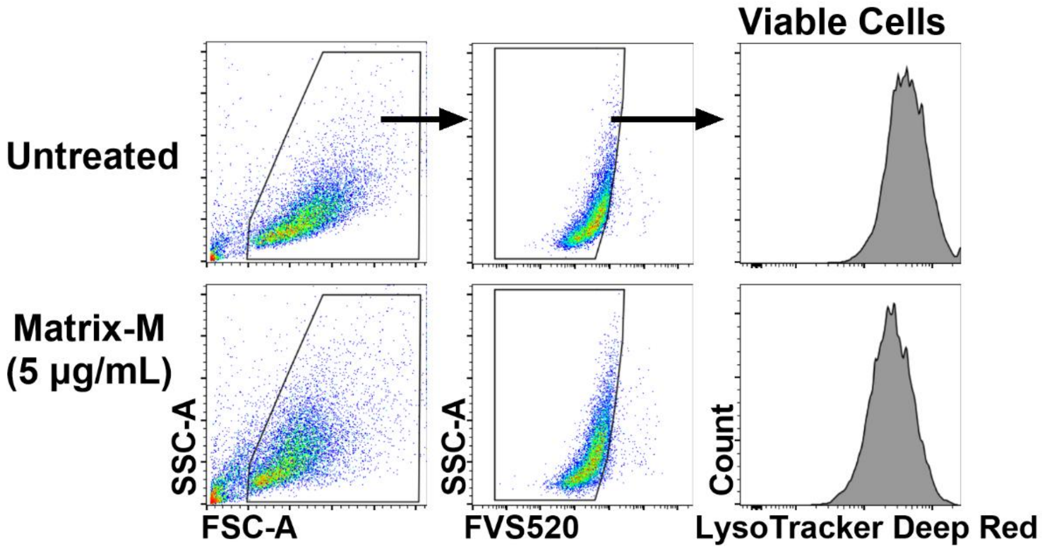


**Supplementary Figure 2:** LMP flow cytometry gating strategy

Lysosomal membrane permeabilization (LMP) in BMDCs after Matrix adjuvant treatment was assessed using a flow cytometry-based assay. A representative gating strategy employed for the flow cytometric analysis to measure LysoTracker Deep Red fluorescence intensity in viable (FVS520 negative) single cells. Gating for doublet cell exclusion was done but is not shown.

**Supplementary Figure 3**

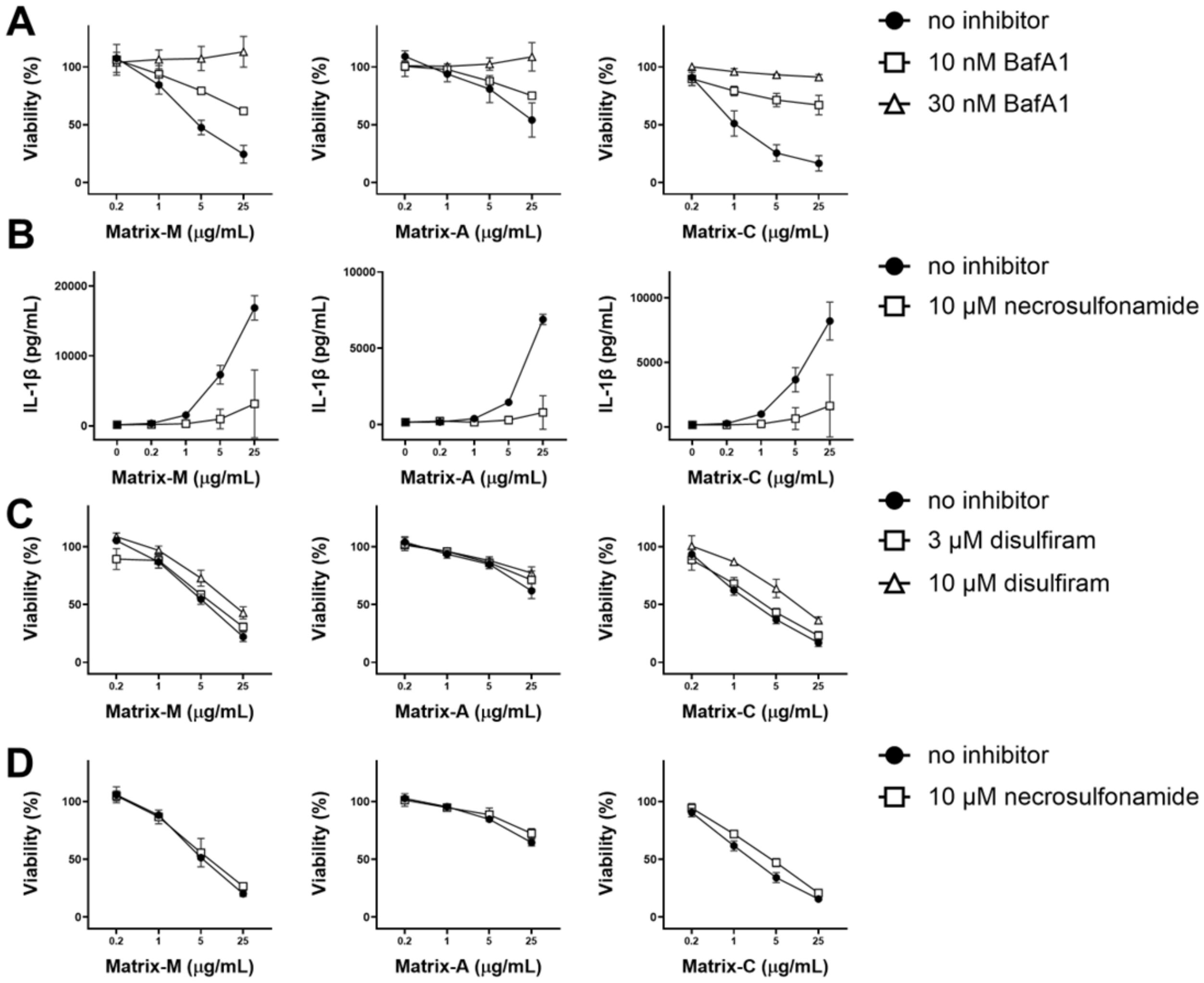


**Supplementary Figure 3**: Viability and IL-1β secretion of Matrix-treated BMDCs

LPS-primed BMDCs were treated with increasing doses of Matrix-M, Matrix-A, or Matrix-C for 5 h. Cell viability was assessed by WST-1 assay (**A, C, D**) and IL-1β levels in supernatants were measured by ELISA (**B**). Before treatment with Matrix adjuvants, cells were pre-treated for 1 h with no inhibitor, V-ATPase inhibitor bafilomycin A1 (**A**), gasdermin D pore formation inhibitors necrosulfonamide (**B, D**), or with disulfiram (**C**) at the indicated concentrations. Results are shown as the mean ± SD from three experiments.

**Supplementary Figure 4**

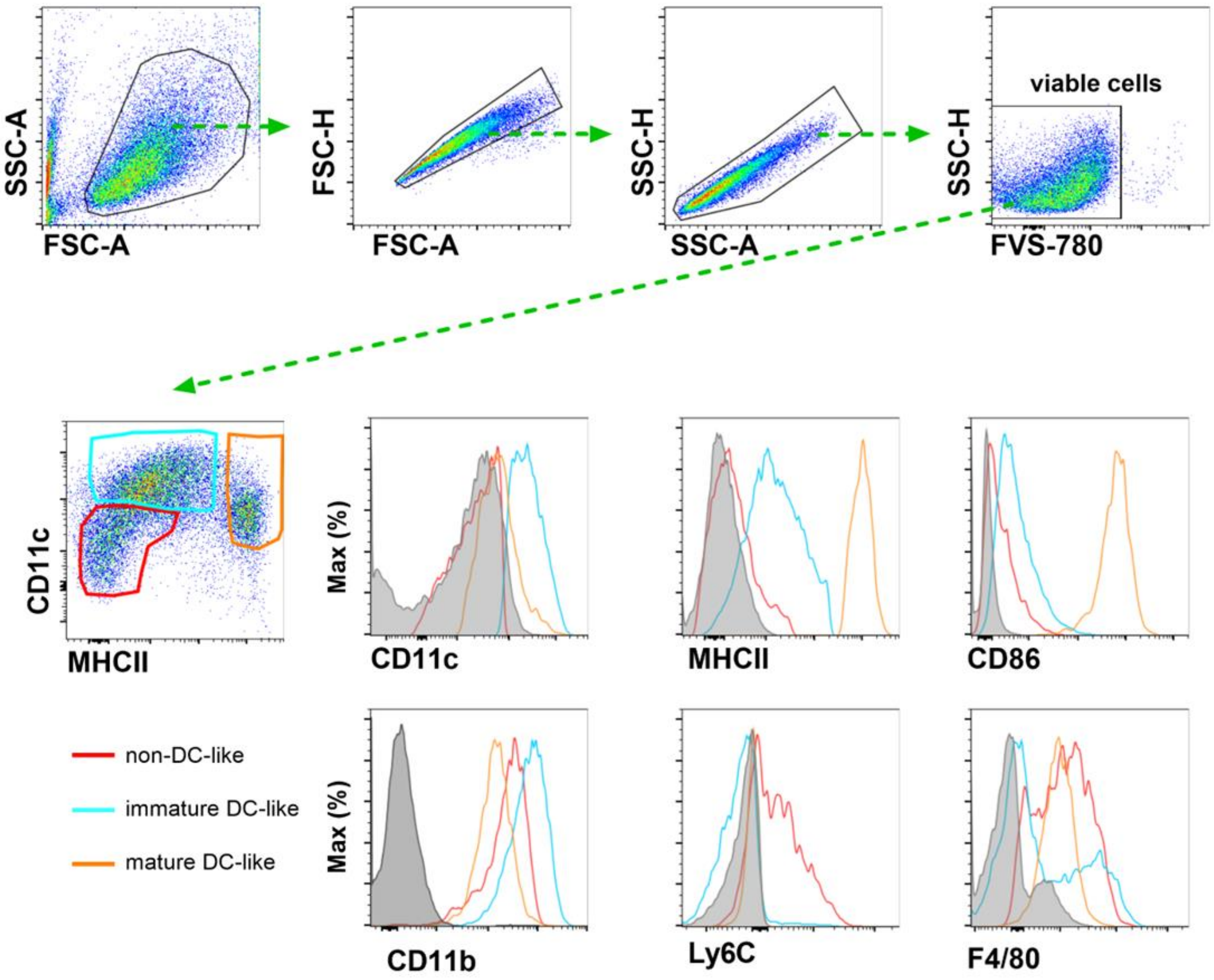


**Supplementary Figure 4:** Gating strategy for BMDC characterization

Gating strategy for BMDCs after 7 days of culture with GM-CSF. The upper panel shows exclusion of cell debris and doublets, further gated for live cells as FVS-780 negative. Live cells were gated as non–DC-like cells (red) that are CD11c- and MHCII-, as immature DC-like (blue) that are CD11c$^{+}$ and negative or low for MHCII, or as mature DC-like cells (orange) that are MHCII high. Histograms show the indicated markers in the three DC subpopulations (red, blue, and orange) and the respective FMO control (gray).

**Supplementary Figure 5**

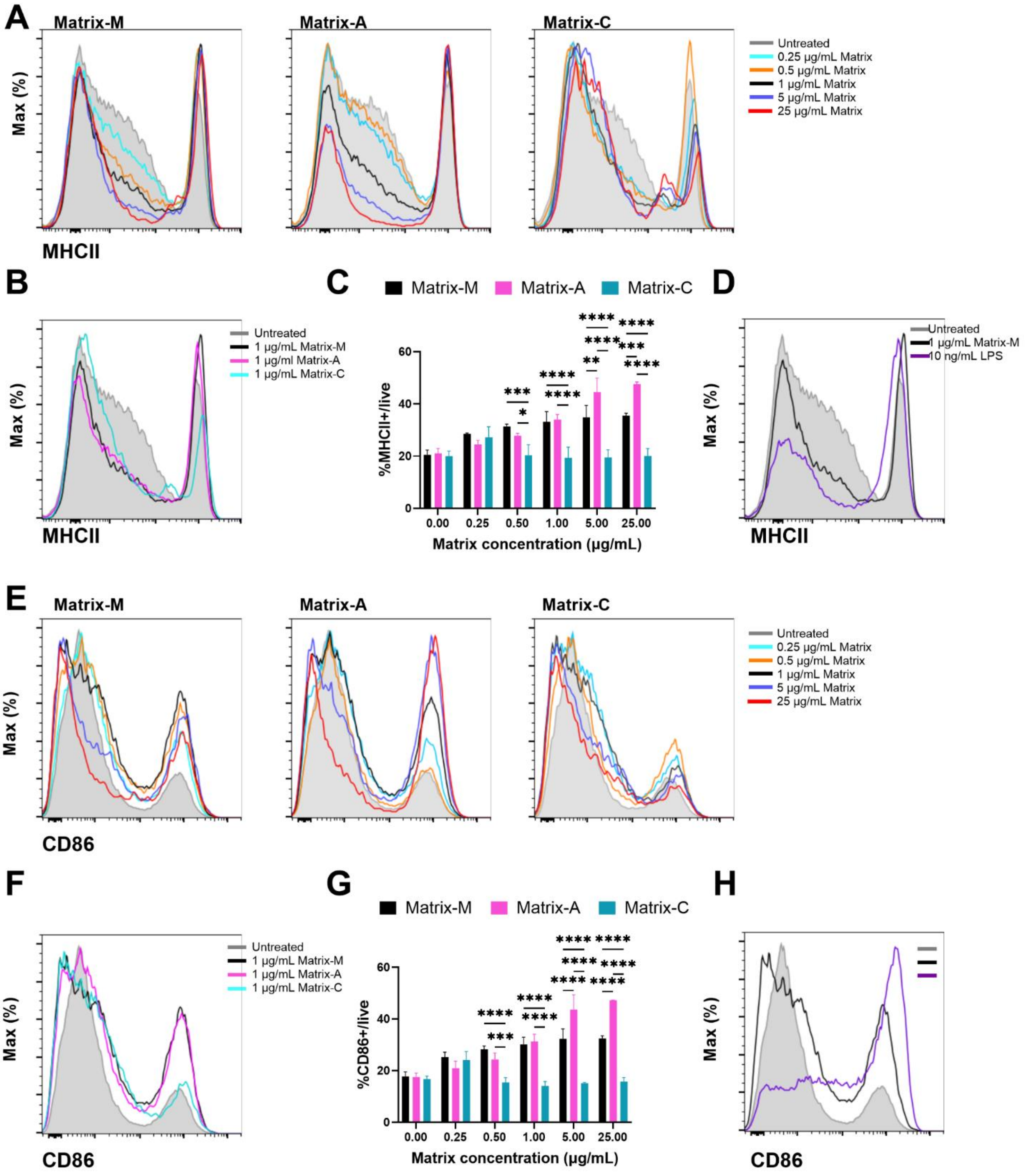


**Supplementary Figure 5:** BMDC activation after Matrix-M, Matrix-A, or Matrix-C treatment

BMDCs were treated for 24 h with the indicated concentrations of Matrix-M, Matrix-A, or Matrix-C and were analyzed by flow cytometry. Representative histograms show the MHCII and CD86 expression levels of live cells for each Matrix treatment at varying concentrations (**A, E**) and for 1 µg/ml Matrix-M, Matrix-A, and Matrix-C (**B, F**). Untreated controls are shown as shaded gray. Summary data showing MHCII and CD86 expression levels in BMDCs treated with Matrix-M, Matrix-A, or Matrix-C (**C, G**). Data represent mean ± SD from three independent experiments. Twoway ANOVA with Tukey's multiple comparisons test was applied for comparison (p-values * for p <0.05, ** p <0.01, *** p<0.001, **** p<0.0001). To compare the effect of 1 µg/ml Matrix-M treatment to LPS (10 ng/ml), a representative histogram shows MHCII (**D**) and CD86 (**H**) expression levels on live BMDCs.

### Supplementary Figure 6

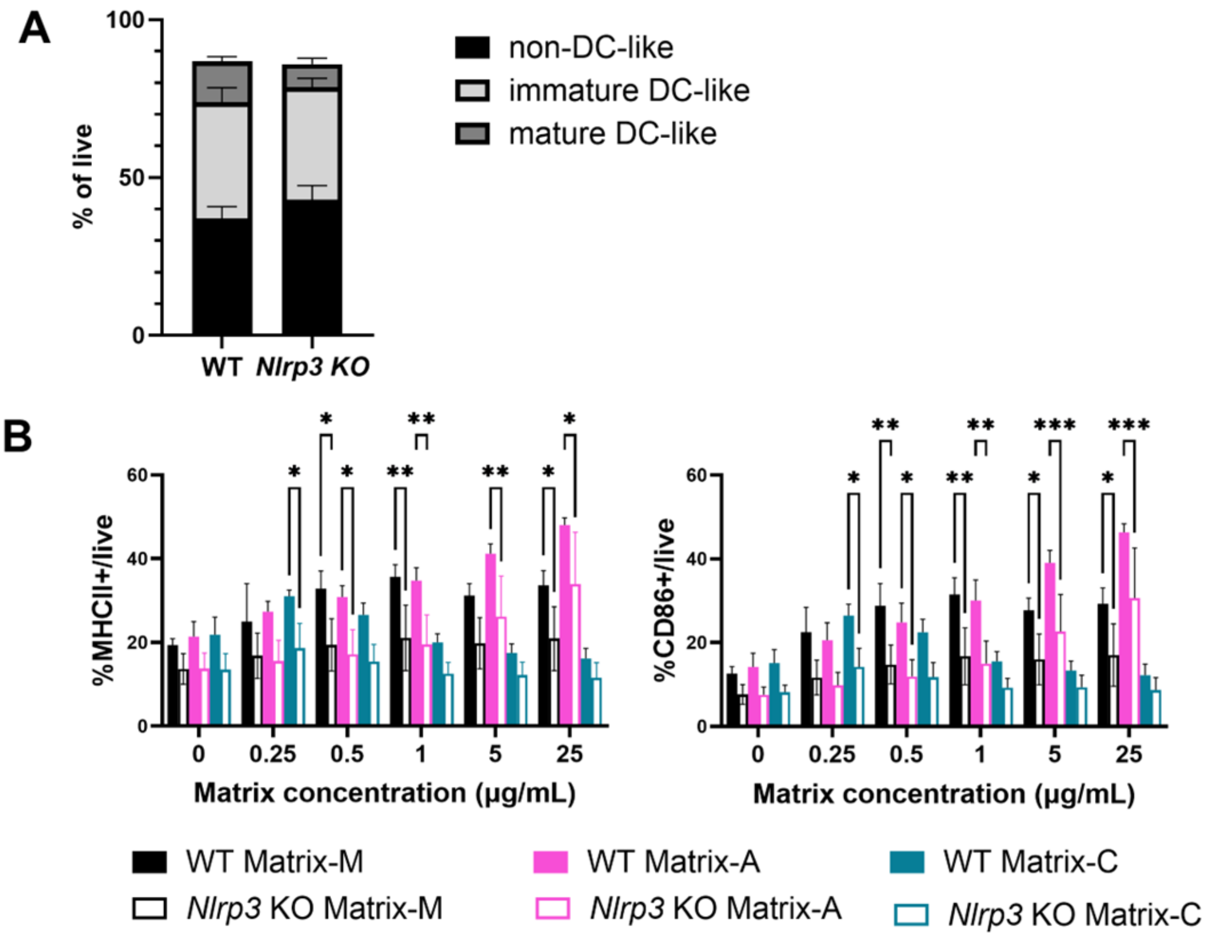


**Supplementary Figure 6:** Phenotype and activation of NLRP3-deficient BMDCs

Untreated BMDCs generated from wild type (WT) and *Nlrp3* KO mice were analyzed by flow cytometry for their phenotype (see also **Supplementary Fig. 4**: non–DC-like, immature DC-like and mature DC-like cells)(**A**) and for their activation based on MHCII and CD86 expression levels (**B**).

**Supplementary Figure 7**

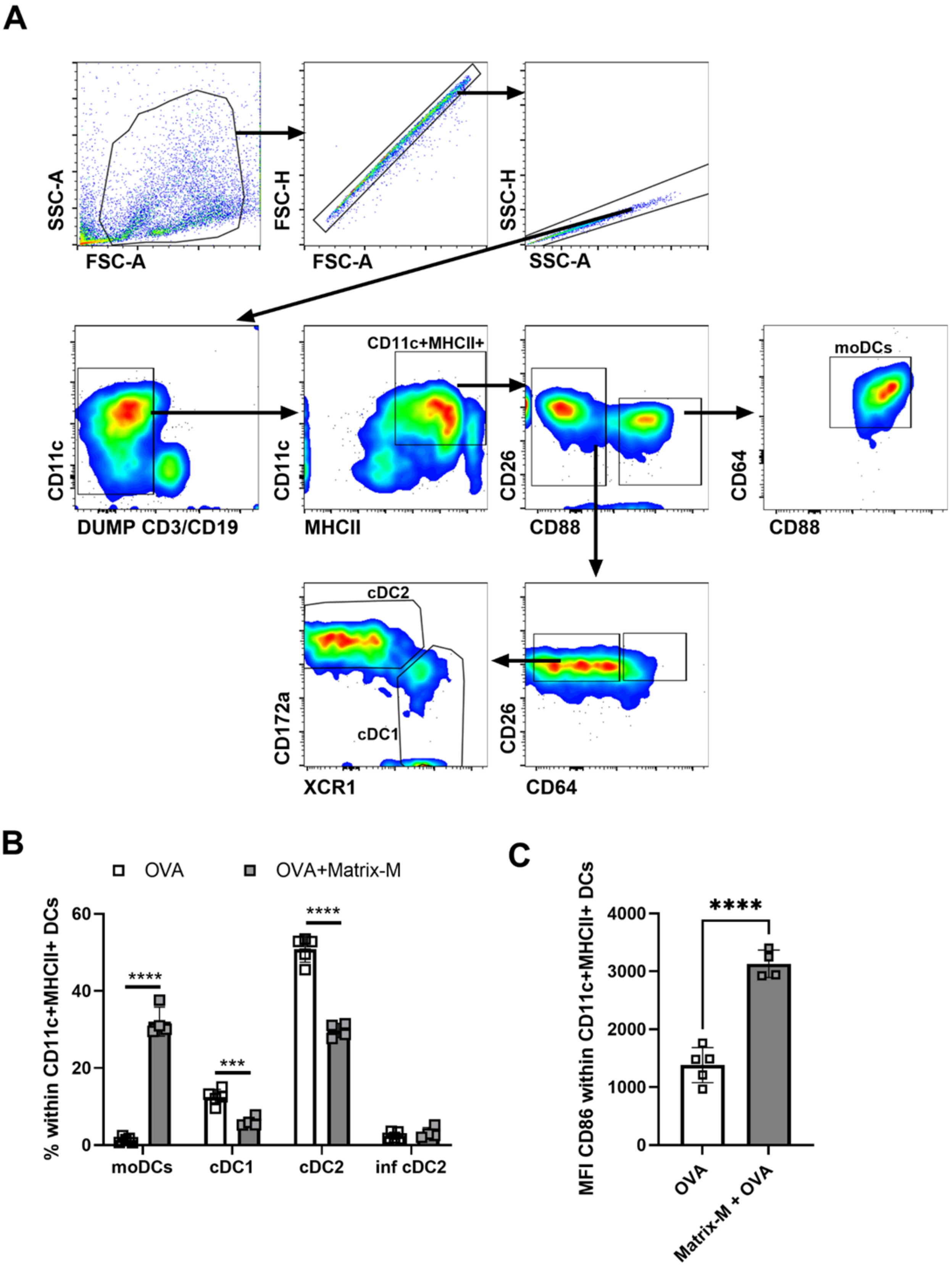


**Supplementary Figure 7:** *Ex vivo* analysis of DCs from immunized mice

C57BL/6 mice were immunized with 10 µg OVA alone or adjuvanted with 5 µg Matrix-M. Iliac lymph node samples were collected 24 h post immunization, processed and sorted for CD11c$^+$ DCs cells. DCs used for T-cell co-culture (Figure 6A–D) were analyzed by flow cytometry with the following gating strategy (**A**): The upper panel shows exclusion of cell debris and doublets and identification of antigen-presenting cells (CD11c$^+$ MHCII$^+$) after excluding lymphocytes (DUMP; CD19$^+$/CD3$^+$ for B and T cells, respectively). Antigen-presenting cells were further classified into CD88$^+$ or CD88$^-$ subsets. Within the CD88- subset, CD26$^+$ cells were identified as cDCs, with CD64$^-$ cDCs subdivided into cDC1 (XCR1$^+$) and cDC2 (CD172a$^+$), while cDCs that were CD64$^+$ were classified as inflammatory cDC2. CD88$^+$ cells expressing CD64 were defined as monocyte-derived DCs (moDCs). The frequencies of these DC subsets from pooled samples (n=4-5) are presented in **(B)**, while the MFI of CD86 within CD11c$^+$MHCII$^+$ DCs is shown in (**C)**. Each symbol represents one individual pool, bar graphs represent mean values, and error bars represent the standard deviation (mean ± SD). Statistical analyses were performed using One-way ANOVA with Bonferroni's multiple comparisons test (B) or an unpaired t-test (C) (*** for p<0.001, **** for p<0.0001).

**Supplementary Figure 8**

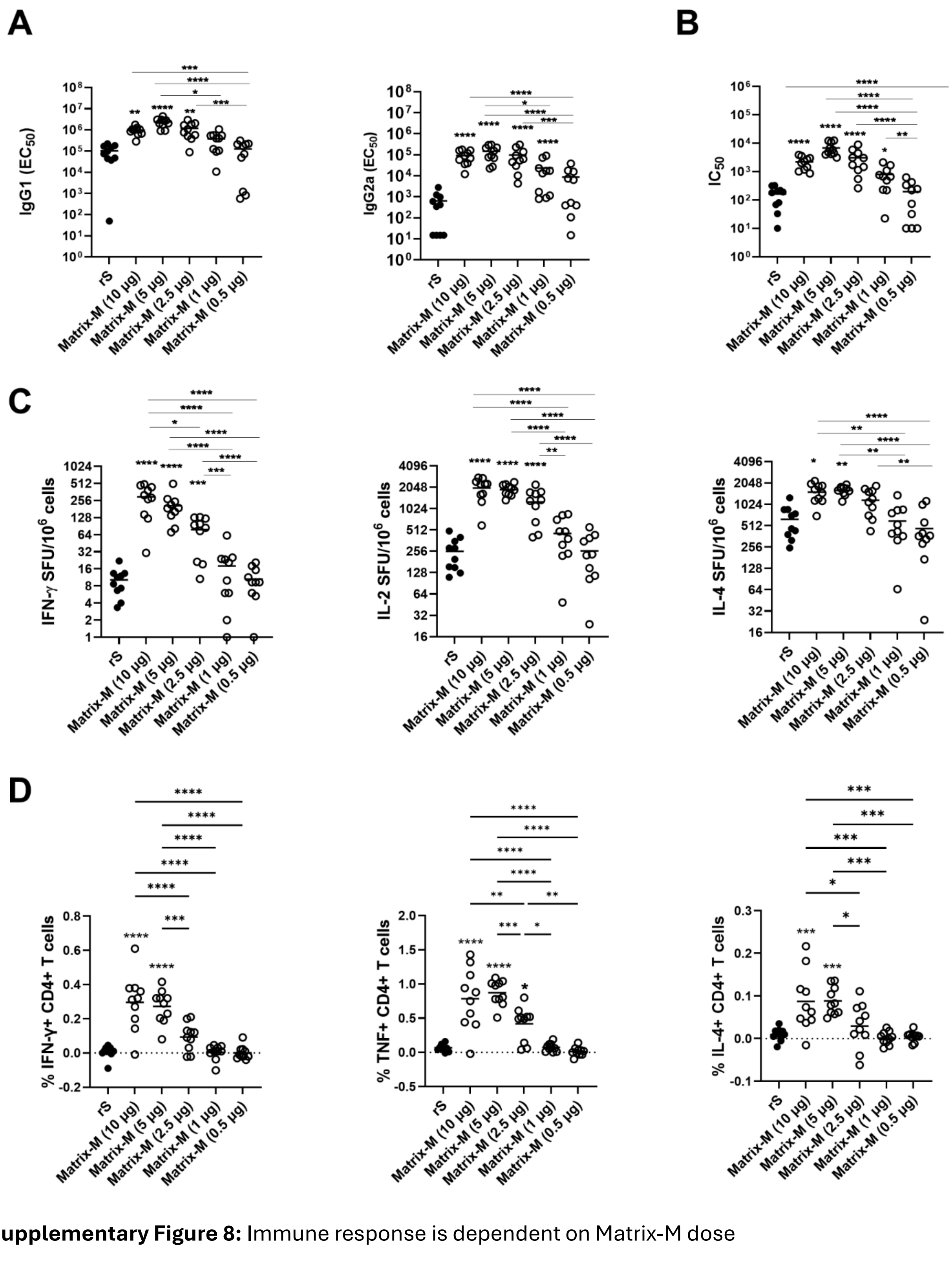


**Supplementary Figure 8:** Immune response is dependent on Matrix-M dose

BALB/c mice (n = 10/group) were immunized with 0.1 µg SARS-CoV-2 rS alone or adjuvanted with the indicated doses of Matrix-M. Serum samples obtained at day 28 (after primary immunization) were evaluated by ELISA for IgG1 and IgG2a (**A**) antibody titers against rS protein and for hACE2 receptor-inhibiting antibody titers (**B**). Splenocytes from individual mice were isolated at day 28 and analyzed for the number of cells producing IFN-γ, IL-2, or IL-4 in response to SARS-CoV-2 rS protein restimulation (**C**). Each symbol represents individual mice, horizontal bars represent geometric mean values. Additionally, splenocytes from individual mice were analyzed by flow cytometry (intracellular staining) for the frequency of $CD4^+$ T cells that are CD44-high and produce IFN-γ, TNF, or IL-4 after rS protein re-stimulations (**D**). Each individual mouse response is shown with each symbol; the solid horizontal bars represent the group mean. The data (**A–D**) were analyzed by one-way ANOVA with Tukey's/Dunnets multiple comparisons test (*for $p<0.05$, ** for $p<0.01$, *** for $p<0.001$, ****for $p<0.0001$). No line indicates a significant difference compared to antigen only, and a line between two groups indicates a significant difference between adjuvanted vaccine groups.
